# KLF4 promotes a KRT13+ hillock-like state in lung squamous cell carcinoma

**DOI:** 10.1101/2025.03.10.641898

**Authors:** Luke T. Izzo, Tony Reyes, Srijan Meesala, Abbie S. Ireland, Lisa Earnest-Noble, Steven Yang, Hari Shankar Sunil, Xiao Chun Cheng, Nomi Tserentsoodol, Sarah B. Hawgood, Carolyn Glass, Edward F. Patz, Benjamin L. Witt, Darren R. Tyson, Kathryn A. O’Donnell, Trudy G. Oliver

## Abstract

Lung squamous cell carcinoma (LUSC) is a basal-like subtype of lung cancer with limited treatment options. While prior studies have identified tumour-propagating cell states in squamous tumours, the broader landscape of intra-tumoural heterogeneity within LUSC remains poorly understood. Here, we employ Sox2-driven mouse models, organoid cultures, and single-cell transcriptomic analyses to uncover previously unrecognized levels of cell fate diversity within LUSC. Specifically, we identify a KRT13^+^ hillock-like population of slower-dividing tumour cells characterized by immunomodulatory gene expression signatures. The tumour hillock-like state is conserved across multiple animal and human-derived models and is present in the majority of human LUSCs as well as head and neck and esophageal squamous tumours. Our findings shed light on the cellular origins of tumour hillock-like states: lung club cells give rise to tumours with luminal hillock-like populations, while basal-like tumour-propagating cells transition into basal hillock-like states, resembling lineage plasticity trajectories of the normal lung. Mechanistically, KLF4 promotes KRT13, a broadly conserved hillock-like state with enrichment of potential therapeutic targets, and resistance to platinum-based chemotherapy. Together, these results provide molecular insights into the lineage plasticity underlying intra-tumoural heterogeneity within LUSC, offering potential avenues for new therapeutic strategies.

## Main

Understanding the cellular complexity of tumours is crucial for improving cancer treatment. Tumours are composed of diverse cell types, including cancer cells, immune cells, and stromal cells, each contributing to disease progression. Among cancer cells, distinct transcriptional states arise not only from genetic alterations but also from non-genetic factors, such as environmental signals and epigenetic modifications^1–3^. Recent advances in single-cell RNA sequencing (scRNA-seq) have transformed our ability to unravel this molecular complexity, revealing previously unappreciated cancer cell states^4–7^. However, the mechanisms driving these transcriptional states in cancer and their impact on therapy resistance remain poorly understood.

Non-small cell lung cancer (NSCLC) represents approximately 85% of all lung cancer cases in the United States^8^. Large-scale genomic sequencing studies have identified genetic alterations commonly found within NSCLC, including lung squamous cell carcinoma (LUSC) and lung adenocarcinoma (LUAD)^9^. While advances in targeted therapies have significantly improved outcomes for LUAD, the lack of actionable targets in LUSC leaves patients with fewer treatment options, underscoring the urgent need for alternative therapeutic strategies for LUSC^10–12^. There is growing evidence that relying solely on cancer genomic alterations for therapy assignment has limitations, which may be due to epigenetic and transcriptional heterogeneity^3,13,14^. Intra-tumoural heterogeneity and the ability of tumour cells to switch between phenotypic states, referred to as plasticity, has recently been highlighted as a hallmark of cancer and a major barrier to cancer treatment^15–17^.

Previous studies investigating intra-tumoural heterogeneity in LUSC identified basal-like tumour-propagating cells (TPCs) that can serially transplant the disease^18–23^. The ability of tumour cell states to mimic normal cell fate lineages (so called “lineage plasticity”) has been observed across various cancer types, including lung cancer, olfactory cancer, and neuroblastoma^24–29^. Basal cells, considered a cell of origin for LUSC^30–32^, are abundant in the major airways where LUSC arises. Basal cells play a central role in maintaining the airway epithelium, responding dynamically to injury and environmental insults such as cigarette smoke, a major risk factor for LUSC^33,34^. Importantly, squamous tumours share key markers and gene expression profiles with basal cells, including the expression of ΔNp63 and KRT5^8^, further supporting the link between basal cells and LUSC pathogenesis. However, whether other cell fates arise from basal-like TPCs in LUSC is largely unexplored.

Lineage-tracing studies show that normal basal cells have the capacity to give rise to multiple differentiated cell types in the adult airway, including neuroendocrine, tuft, ionocyte, club, ciliated, and newly-identified hillock cells^6,7,35^. Hillock cells were first reported in single cell studies that suggested basal cells can pass through a hillock state before progressing toward a club fate^6^. Normal hillock cells have been identified in multiple tissues including cartilaginous airways of the lung and in the prostate^36^, and they are marked by KRT13 expression with signatures of immunomodulation and squamous differentiation^6,7^. Recent work stratified hillock cells into two subsets, proliferative basal hillock cells and protective luminal hillock cells^37^. Basal hillock cells have a remarkable ability to repopulate the tracheal epithelium after injury. Luminal hillock cells are morphologically and transcriptionally similar to differentiated squamous cells and maintain basal-hillock cell integrity through resistance to a myriad of injuries^37^. The function of normal hillock cells is only beginning to be understood, and whether they have a transcriptional or functional counterpart in cancer remains completely unexplored to our knowledge. Understanding whether hillock-like cells exist in squamous tumours could provide new insights into the transcriptional plasticity driving LUSC progression and resistance to therapy.

Multiple genetically-engineered mouse models (GEMMs) of LUSC have been developed^38–43^. While the basal cell is thought to be the most physiologically relevant cell-of-origin for LUSC, multiple cell types have been shown to give rise to LUSC^30,38^. A key difference between mouse and human lungs is that basal cells are largely confined to the trachea in the mouse, and most GEMMs rely on delivery of virus to more distal regions of the lung where squamous tumours arise via transdifferentiation^40,44–48^. Importantly, mouse squamous tumours that arise via transdifferentiation highly resemble their human tumour counterparts both molecularly and histologically^18,30,40,49–51^. Due to this transcriptional and histopathological resemblance, mouse models provide useful tools for interrogating intra-tumoural heterogeneity and the molecular mechanisms that regulate cell state in squamous tumours.

Kruppel-like factor 4 (KLF4) is a transcription factor that plays an essential role in proliferation, differentiation, and stem cell pluripotency in a context-specific manner^52–54^. In both the skin and esophageal epithelium, KLF4 is restricted to the suprabasal layer and is necessary for differentiation of specific cell types^55–58^. Apart from its role in normal development, KLF4 has been implicated in multiple cancers—able to adopt either an oncogenic or a tumour suppressive role in a context-dependent manner^52^. Despite KLF4’s link to normal squamous differentiation, the function of KLF4 in LUSC and in the hillock cell state has not been explored. Here, by investigating the cellular origins and transcriptional regulation of the KRT13^+^ hillock-like state in LUSC, this study aims to uncover fundamental mechanisms of tumour heterogeneity, paving the way for novel therapeutic strategies targeting transcriptional plasticity in lung squamous cell carcinoma.

## Results

### KRT13 is highly enriched in lung squamous cell carcinoma

Recent single cell studies have identified novel lung cell populations^6,7,37^, including KRT13^+^ hillock cells. To determine whether a similar state exists in lung cancer models, we queried *Krt13* expression across multiple lung cancer GEMMs representing the three major histological subtypes of lung cancer: adenocarcinoma (LUAD), squamous (LUSC), and small cell lung cancer (SCLC). We collected multiple samples of adenocarcinoma from the lungs of *Kras^G12D/+^;Trp53^fl/fl^*(KP) mice, squamous tumours from *Sox2^LSL/LSL^;Nkx2-1^fl/fl^;Lkb1^fl/fl^*(SNL) or *Lkb1^fl/fl^;Pten^fl/fl^* (LP) mice, and SCLC from *Rb1^fl/fl^;Trp53^fl/fl^;Rbl2^fl/fl^* (RPR2) or *Rb1^fl/fl^;Trp53^fl/fl^;Myc^T58A/T58A^* (RPM) mice and performed RNA-seq^18,40,49,59–61^. We observed abundant *Krt13* expression restricted specifically to the squamous GEMMs (**Fig. 1a**). To test whether this association was consistent at the protein level, we performed immunohistochemistry (IHC) on tumour-bearing lung tissue from the lung cancer GEMMs. KRT13 expression was highly enriched in squamous but not in adenocarcinoma or SCLC tumours (**Fig. 1b,c**). To determine whether this association was consistent in human squamous tumours, we examined RNA-seq data derived from human lung adenocarcinoma and squamous tumours from The Cancer Genome Atlas (TCGA). Compared to lung adenocarcinoma, we observed significantly greater *KRT13* expression in human squamous tumours (**Fig. 1d**). Tissue microarrays of human lung cancer (n = 57 total samples, n = 41 containing tumour tissue) stained for KRT13 also had significantly increased expression in human squamous compared to adenocarcinoma samples, with the majority of squamous patient biopsies displaying KRT13 positive cells (16 of 20) (**Fig. 1e,f**). Notably, both mouse and human squamous tumours exhibited intra-tumoural heterogeneity in KRT13 expression (**Fig. 1b,e**). Overall, these data suggest KRT13^+^ cells are highly enriched in subpopulations of mouse and human squamous tumours.

**Figure 1.**
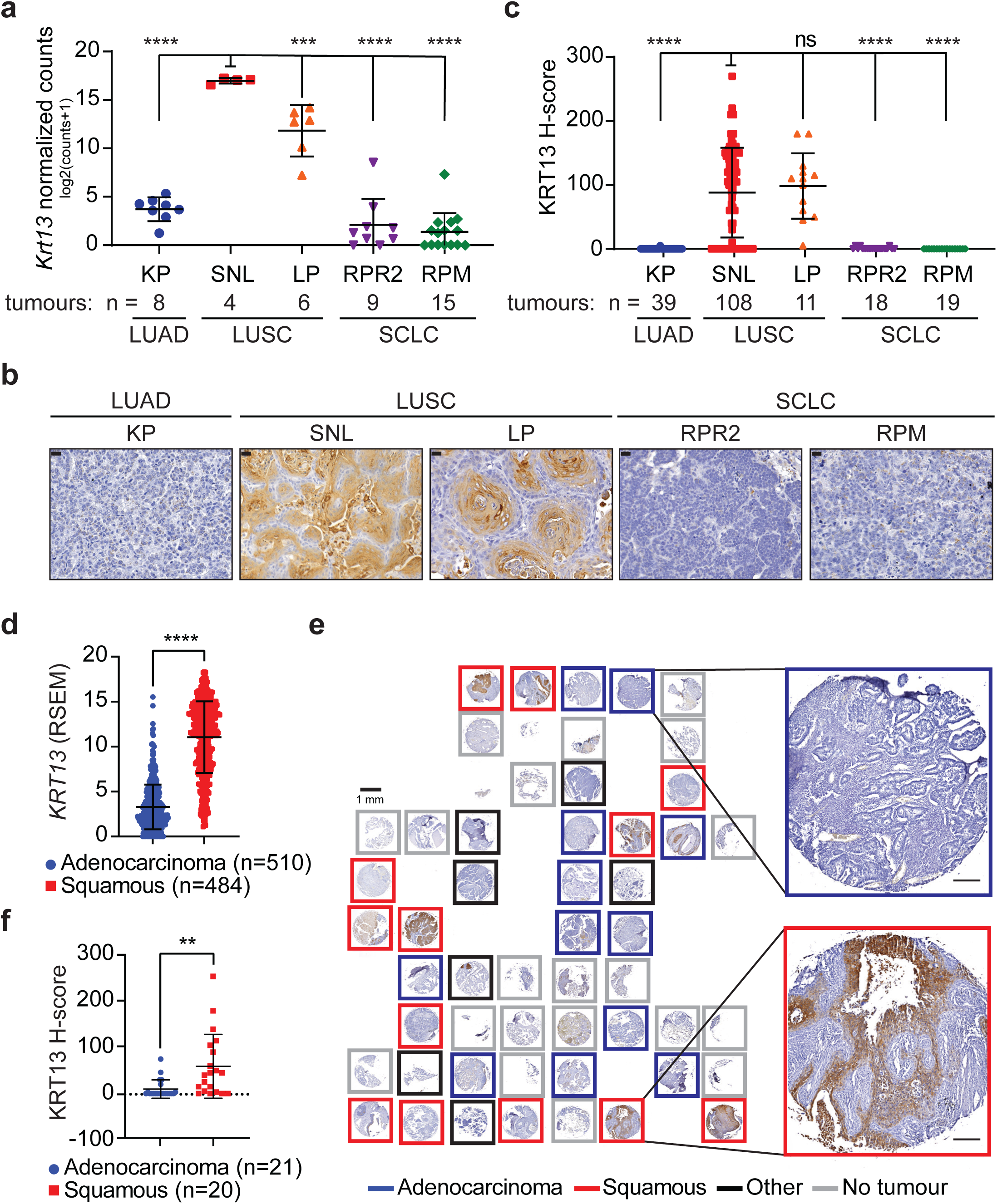
KRT13 is highly enriched in lung squamous cell carcinoma. **a)** *Krt13* expression shown as log2 normalized counts by RNA-seq from lung tumours in indicated GEMMs grouped according to histological type. Tumours were collected from the following number of mice for each genotype: RPR2 (n=8), RPM (n=12), SNL (n=4), LP (n=5), KP (n=6). Statistical significance determined by one-way analysis of variance (ANOVA) and p-values are for all pairwise comparisons to SNL. Lung squamous cell carcinoma (LUSC); lung adenocarcinoma (LUAD); small cell lung cancer (SCLC). *Sox2^LSL/LSL^;Nkx2-1^fl/fl^;Lkb1^fl/fl^*(SNL); *Lkb1^fl/fl^;Pten^fl/fl^* (LP); *Rb1^fl/fl^;Trp53^fl/fl^;Myc^T58A/T58A^*(RPM); *Rb1^fl/fl^;Trp53^fl/fl^;Rbl2^fl/fl^* (RPR2); *Kras^G12D/+^;Trp53^fl/fl^* (KP). **b)** Representative immunohistochemistry (IHC) for KRT13 in indicated GEMMs. Scale bar, 20 µm. **c)** H-score quantification of IHC in **1b** (see **Methods**). Number of tumours indicated in figure. Statistical significance was determined by one-way ANOVA and p-values are for all pairwise comparisons to SNL. **d)** *KRT13* expression shown as RSEM from human lung adenocarcinoma and squamous cell carcinoma from TCGA. Number of tumours indicated in figure. Two-tailed t-test. **e)** Representative (1 of 2 TMA slides) KRT13 IHC in human lung tumour tissue microarrays (red = squamous cell carcinoma; blue = adenocarcinoma; black = NSCLC/large cell, grey = section without tumour tissue or not enough material to score). Scale bar, 1 mm. Higher magnification insets for adenocarcinoma (blue) and squamous cell carcinoma (red); scale bar, 100 µm. **f)** H-score quantification of IHC in **1e** with number of tumours indicated. Two-tailed t-test. **For all panels**, each point represents an individual tumour, and error bars represent mean +/-SD. ns = not significant; **p ≤ 0.01; ***p ≤ 0.001; ****p ≤ 0.0001.

### KRT13 is expressed in a non-basal subpopulation of LUSC

Previous studies have shown functional heterogeneity in LUSC, including a TPC population marked by NGFR^+^SCA1^+^ expression with the capacity to serially transplant tumours^18^. Squamous tumours in most GEM models have been shown to arise via transdifferentiation from a mucinous-like state^9,40,44–46^. Due to potential transcriptional and functional heterogeneity, and the heterogeneous expression of KRT13 in squamous tumours (**Fig. 1b,e**), we sought to assess *Krt13* expression at the single cell level in squamous GEMMs via single cell RNA-sequencing (scRNA-seq). We flow-sorted CD45-negative (CD45^-^) GFP-positive (GFP^+^) tumour cells from Adenoviral-CMV-Cre-infected (Ad-CMV-Cre) SNL mice utilizing the GFP reporter in the *Sox2-Ires-Gfp* allele. Following sorting, CD45^-^ GFP^+^ tumour cells were subject to scRNA-seq (see **Methods**). We obtained 3,663 tumour cells, but very few of these cells expressed canonical basal-associated squamous markers *Krt5*, *Trp63*, *Pitx1* and *Ngfr* that are known to be present in squamous tumours by IHC. Rather, the majority of CD45^-^GFP^+^ tumour cells expressed markers of mucinous adenocarcinoma. To capture more basal-like squamous tumour cells, we flow-sorted Ad-CMV-Cre-infected SNL tumours and isolated CD45^-^GFP^+^NGFR^+^ cells (**Extended Data Fig. 1a**), which are reported to include the TPC population. We obtained 4,363 additional tumour cells via scRNA-seq that were enriched for squamous marker genes (**Fig. 2a-c**). Consistent with previous studies^40,62^, mucinous markers *Spdef, Hnf4a*, and *Gkn1* were low in tumour cells that expressed basal-associated squamous markers (**Fig. 2c**). *Krt13* expression was overall low, but present in tumour cell clusters expressing basal markers and absent in cells expressing mucinous adenocarcinoma markers (**Fig. 2d**). Consistently, IHC on serial sections from Ad-CMV-Cre-infected SNL tumours showed that KRT13 was enriched in KRT5^+^ squamous regions and absent in HNF4A^+^ mucinous regions (**Fig. 2e**). Co-staining for KRT13 and basal marker TRP63 showed that KRT13 was highly expressed in cells that were adjacent to, but distinct from, TRP63-expressing cells (**Fig. 2f**). Together, these data suggest that KRT13^+^ cells are enriched in a subpopulation of non-mucinous, non-basal squamous tumour cells.

**Figure 2.**
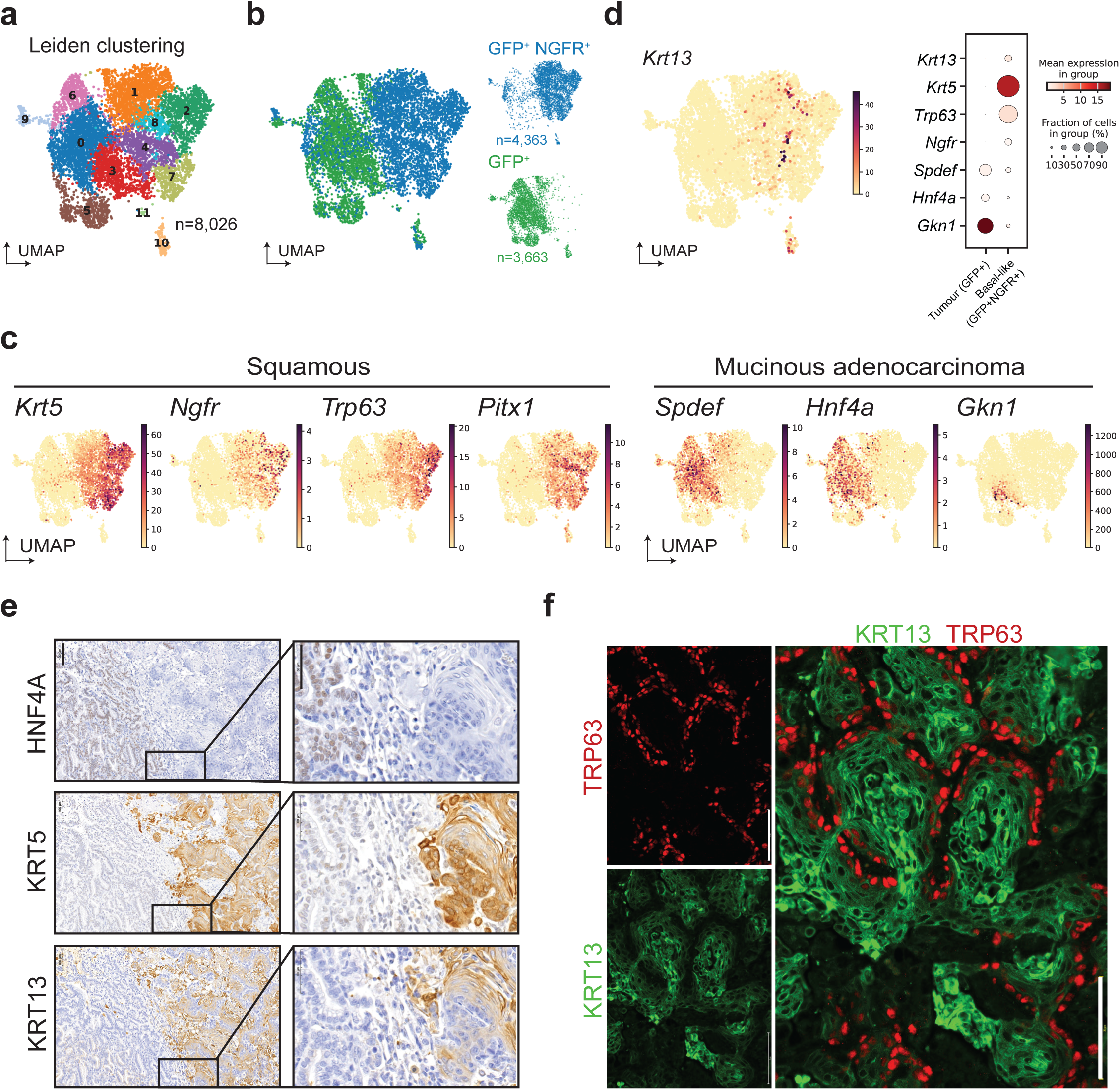
KRT13 is expressed in a non-basal subpopulation of LUSC. **a)** ScRNA-seq of Ad-CMV-SNL tumour cells labeled by Leiden clustering in UMAP space. **b)** Ad-CMV-SNL tumour cells labeled by sorted population (GFP^+^ or GFP^+^/NGFR^+^) in UMAP space as in **2a**. **c)** Expression of indicated squamous or mucinous adenocarcinoma gene markers in UMAP space from cells in **2a**. **d)** ScRNA-seq expression of *Krt13* projected on UMAP space from cells in **2a** and dotplot of gene expression from scRNA-seq separated by flow sort. Dot size indicates percent of cells expressing each transcript and the color indicates the average expression per cluster indicated in the figure. **e)** Representative IHC from serial sections in Ad-CMV-SNL-tumour-bearing lungs harbouring both squamous cell carcinoma and mucinous adenocarcinoma. Representative of n = 6 Ad-CMV-SNL mice. Scale bar, 100 µm for left column; 50 µm for high-magnification insets on right column. **f)** Representative immunofluorescence co-staining for KRT13 (green) and TRP63 (red) in Ad-CMV-SNL tumours. Representative of n = 4 Ad-CMV-SNL mice. Scale bars, 75 µm. **See also Extended Data Fig. 1.**

### Club-initiated tumours have early and enriched KRT13 expression

Given that normal *Krt13*^+^ lung cells can act as an intermediate state between basal and club cells^6^, and club cells can regenerate basal cell populations^63^, we reasoned that club-initiated tumour cells may have the capacity to dedifferentiate to a KRT13^+^ state. Using adenoviral delivery of Cre controlled by a club cell-specific promoter (CCSP), we induced tumours from club cells in SNL mice to determine whether this would alter the relative abundance or quality of *Krt13^+^* tumour cell populations by scRNA-seq. Similar to our approach with Ad-CMV-Cre-initiated tumours, we first sorted Adenoviral-CCSP-Cre-driven (Ad-CCSP-Cre) CD45^-^GFP^+^ tumour cells (CCSP-GFP^+^, “S1”) and found a lack of basal-associated squamous marker gene expression. We subsequently performed a second round of sequencing from Ad-CCSP-Cre infected SNL lungs, containing a mixture of sorted CD45^-^GFP^+^ (CCSP-GFP^+^, “S2”) and CD45^-^GFP^+^NGFR^+^ basal-enriched tumour cells (see **Methods**). For subsequent scRNA-seq analysis, cells from both collections were combined with cells derived from Ad-CMV-Cre-infected tumours. Following pre-processing, batch correction, and removal of normal lung and immune cell populations (see **Methods**), we obtained 4,012 club cell-derived tumour cells that largely clustered distinctly from Ad-CMV-initiated populations (**Fig. 3a,b**). Sorting based on NGFR expression enriched for cells expressing basal markers genes (**Extended Data Fig. 2a**). We confirmed the presence of both mucinous adenocarcinoma and squamous populations in Ad-CCSP-derived tumour cells (**Fig. 3b,c**), demonstrating that club cells can act as a cell of origin for squamous tumours in the SNL model. *Krt13*^+^ cells were highly enriched in Ad-CCSP- versus Ad-CMV-initiated tumour cells, particularly in clusters that lacked mucinous and basal squamous markers (**Fig. 3b-d**). We assessed expression of another gene, *Tmprss11b*, that correlates with poor survival in NSCLC^64,65^. *Tmprss11b* is enriched in a similar population of cells as those expressing *Krt13* and correlated with *Krt13* expression (**Extended Data Fig. 2b,c**). Interestingly, in contrast to Ad-CMV-driven tumours, KRT13 was present in mucinous adenocarcinoma as well as in squamous histopathologies in Ad-CCSP-initiated tumours (**Fig. 3e,f**). Given that prior studies have shown that mucinous adenocarcinoma precedes squamous differentiation^40,46,47^, these data suggest KRT13 is enriched and may emerge earlier in club-initiated tumours compared to Ad-CMV-initiated tumours.

**Figure 3.**
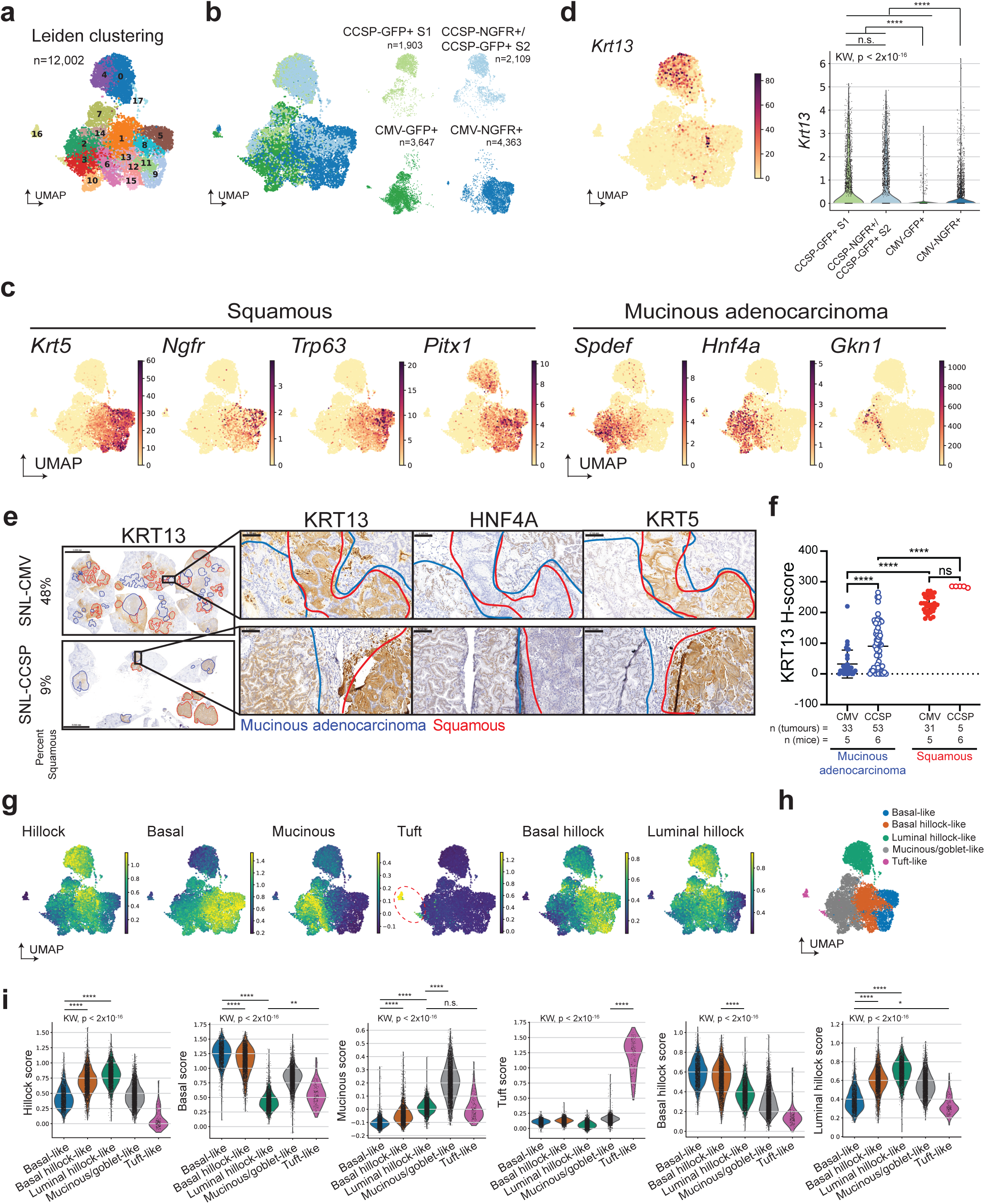
Club-initiated tumours have early and enriched KRT13 expression. **a)** ScRNA-seq UMAP of primary Ad-CMV- and Ad-CCSP-SNL tumour cells labeled by Leiden clustering. **b)** Ad-CMV- and Ad-CCSP-initiated SNL tumour cells labeled by sorted population (GFP^+^ or GFP^+^/NGFR^+^) in UMAP space as in **3a**. **c)** Expression of indicated squamous or mucinous adenocarcinoma gene markers in UMAP space from cells in **3a**. **d)** scRNA-seq expression of *Krt13* projected on UMAP space from cells in **3a** and violin plots for *Krt13* expression by sample. Kruskal-Wallis followed by Dunn’s test with Bonferroni correction, not all comparisons shown. **e)** Representative KRT5, HNF4A, and KRT13 IHC for Ad-CMV- and Ad-CCSP-initiated SNL tumours. Squamous (red) and mucinous adenocarcinoma (blue) tumour lesions are outlined. Scale bar, 5 mm for left panels; 2 mm for high-magnification insets on serial sections in right three panels. Representative of n = 5 Ad-CMV-SNL and n = 6 Ad-CCSP-SNL mice. **f)** H-score quantification of IHC in **3e**. Two-way ANOVA with a Sidak correction for multiple comparisons. Not all comparisons are shown. Each point represents an individual tumour, and error bars represent mean +/- SD. ns = not significant; ****p ≤ 0.0001. **g)** Gene scores applied to SNL tumour cells projected onto UMAP space from **3a** (see **Methods** and **Supplementary Tables 1-5**). **h)** Cell type classification based on gene and gene signature score expression from cells in **3a**. **i)** Violin plots of gene score expression by cell type assignment. Kruskal-Wallis followed by Dunn’s test with Bonferroni correction, all comparison p < 0.0001 unless otherwise shown. **See also Extended Data Fig. 2 and 3.**

### KRT13 marks a hillock-like state in squamous tumours

To determine whether *Krt13^+^* cells exhibit a hillock-like phenotype in LUSC, we generated a “hillock-cell signature” by identifying normal hillock-cell specific genes that were conserved across two independent datasets^6,7^ (see **Methods** and **Supplementary Table 1**). We applied the hillock-cell signature to the mouse tumour scRNA-seq data. Tumour cells with a high hillock signature strongly overlapped with *Krt13*-expressing cells and were present in both Ad-CMV and Ad-CCSP-initiated tumours (**Fig. 3d,g**). Additionally, tumour cells with a high hillock score were largely distinct from cells with the highest mucinous and basal cell signatures (**Fig. 3g**) (see **Methods, Supplementary Tables 2 and 3**), suggesting that *Krt13* marks a transcriptionally distinct hillock-like state in lung squamous cell carcinoma, as it does in the normal airway epithelium. To query potential similarities between normal lung and airway cell lineages and tumour heterogeneity, we applied additional normal cell type signatures to the scRNA-seq data. Consistent with known relationships between goblet cell phenotypes and mucinous adenocarcinomas, these two signatures largely overlap and cluster together by correlation analysis (**Fig. 3g** and **Extended Data Fig. 2d, e**). The club cell signature overlaps most with the hillock signature and partially overlaps with the basal signature (**Extended Data Fig. 2d,e**) in line with known lineage relationships between normal club, hillock, and basal cells^6^. Next, we generated gene sets derived from a recent study^37^ to distinguish normal basal hillock and luminal hillock cells from other cancer states (see **Methods**, **Supplementary Tables 4 and 5**). Applying these signatures to the mouse tumour scRNA-seq dataset revealed that *Krt13^+^* hillock-like tumour cells overlapped most with the luminal hillock score, while the basal hillock score was highest in the basal-like cells (**Fig. 3g** and **Extended Data Fig. 2e**). Together, these data suggest that *Krt13* is enriched in hillock-like cells that may mark both luminal or basal hillock-like populations, which is influenced by cell of origin.

### Luminal hillock-like cells have reduced proliferation and express immunomodulatory gene signatures

Next, using marker gene expression and gene signature scores of normal lung and airway cell types, we assigned cell states to Leiden clusters in the mouse tumour scRNA-seq dataset. Tumour cells were assigned to either basal-, basal hillock-, luminal hillock-, or mucinous/goblet-like states (**Fig. 3g-i** and **Extended Data Fig. 3a–d**). In addition, a small population of tumour cells with a tuft cell gene expression program was identified that clustered separately from other cell populations (**Fig. 3g-i**, **Extended Data Fig. 2e**, and **Extended Data Fig. 3a–c,** see **Methods**)^6^. Recent reports have noted tuft-like tumour cells in or near squamous tumours^66–68^. Consistently, our lab recently discovered that basal cells can simultaneously give rise to tuft-like SCLC and squamous tumours under specific genetic conditions, and combined SCLC in patients can harbour both tuft-like SCLC and squamous regions^68,69^, suggesting a transcriptional relationship between tuft and basal states in lung cancer. These tuft-like cancer cells were enriched in Ad-CMV- compared to Ad-CCSP-driven tumours, suggesting a potential influence of cell of origin (**Extended Data Fig. 3e**).

Ad-CCSP-driven tumours contained the majority of luminal-hillock-like cells, whereas Ad-CMV-driven tumours contained the majority of basal-hillock-like cells (**Extended Data Fig. 3e**). Notably, luminal hillock-like tumour cells exhibit intermediate club, goblet, and mucinous signatures compared to the mucinous and basal-like populations (**Fig. 3g-i** and **Extended Data Fig. 3d**). Together, these data suggest that cell of origin impacts the presence of different hillock-like states and that tumour lineage plasticity in SNL squamous tumours may mirror normal developmental programs, i.e. transcriptional relationships between club, hillock, and basal cell states^6,37^ (**Extended Data Fig. 3f**).

To uncover transcriptional patterns associated with each cell state in an unbiased manner, we identified top differentially-expressed genes (DEGs) in the assigned luminal-hillock, basal-hillock, or basal-like clusters comparing each cell state to all other cells and performed ENRICHR pathway analysis (**Supplementary Table 6**). Luminal hillock-like cells were enriched for signatures of immune pathways, neutrophil degranulation, and cytokine signaling (**Fig. 4a**), with genes driving the signatures including *B2m, Lgals3, and Il1a* (**Supplementary Table 7**). We also assessed gene expression related to immunomodulation including interferon signaling, immune checkpoint and antigen presentation (**Extended Data Fig. 4a**). Luminal hillock-like cells had notable enrichment for interferon regulatory gene expression (*Irf1*, *Irf7*, and *Ifr9*) and immune checkpoint genes including *Cd274* (encoding PD-L1), *Ceacam1*, and *Vtcn1*. Both basal and basal hillock-like clusters were enriched for proliferation-related gene signatures such as translation and metabolic processes (**Fig. 4b**), suggesting luminal-hillock-like cells may be less proliferative compared to basal and basal-hillock-like tumour cells. Consistently, KI67 expression was restricted to TRP63^+^KRT13^-^ cells and nearly absent from KRT13^+^ regions within squamous tumours by immunostaining (**Fig. 4c**). Additionally, cell cycle-specific gene signatures in scRNA-seq data revealed that the luminal-hillock-cell state has a lower proportion of cells in S and G2/M phases compared to basal and basal-hillock states, consistent with decreased proliferation (**Extended Data Fig. 4b**). Basal-like and basal-hillock-like states have similar gene expression compared to the remaining cells in our data (**Fig. 4b** and **Supplementary Table 6**). To determine differences between the populations, we directly compared the basal-like and basal hillock-like cell states. This analysis revealed that basal hillock-like cells have elevated expression of immunomodulatory signatures and increased expression of tight junction genes compared to basal-like cells, whereas basal-like cells are enriched for translation-related signatures (**Extended Data Fig. 4c-d** and **Supplementary Table 6**). Altogether these data suggest that the KRT13^+^ luminal-hillock-like state has relatively lower proliferation than basal-like populations and that the basal-hillock-like population shares characteristics of the basal- and luminal-hillock-like state.

**Figure 4.**
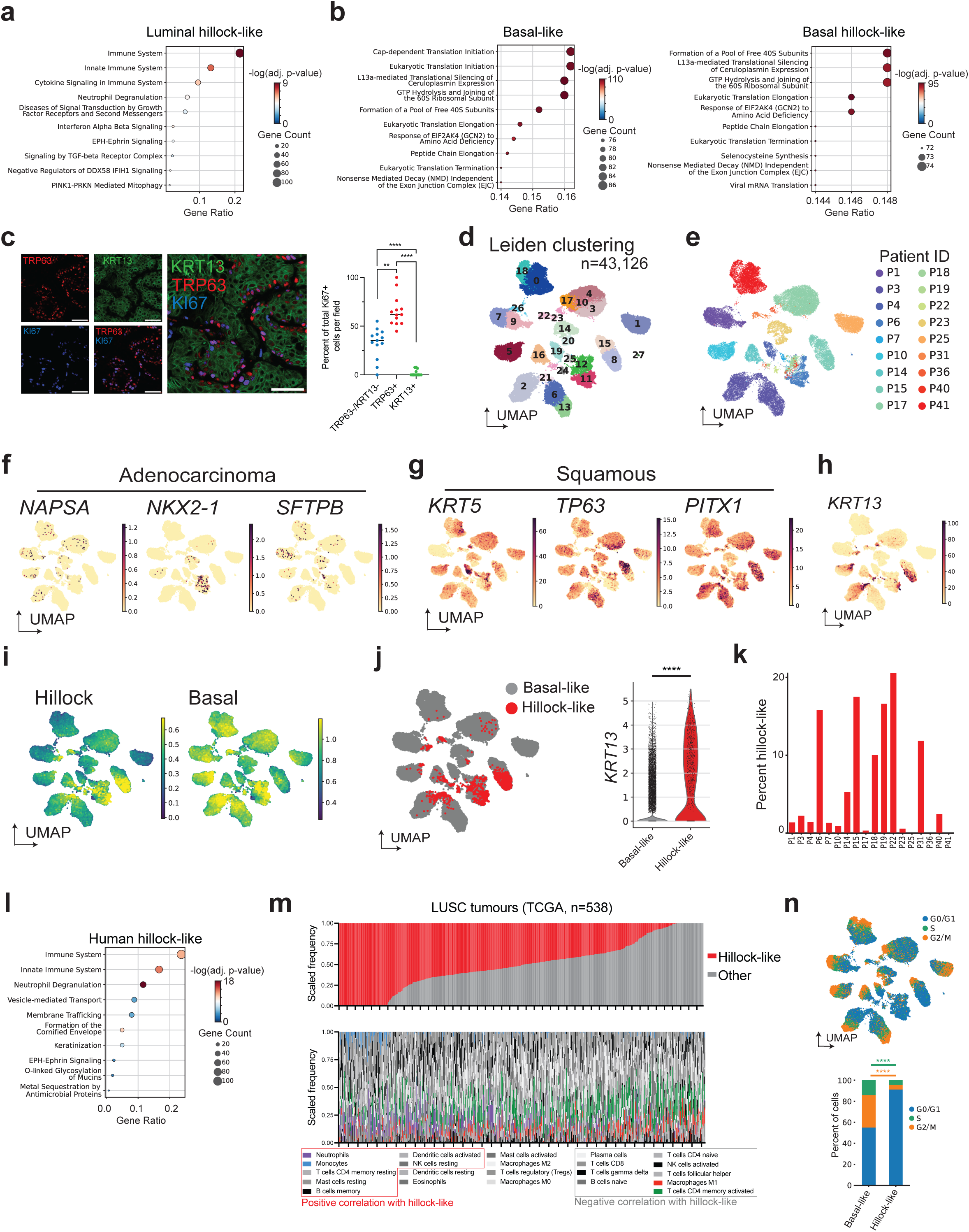
KRT13 marks a hillock-like state in squamous tumours. **a,b)** Gene set enrichment analysis comparing SNL tumour cells in **3a** by assigned cell type. The ENRICHR gene set used was ‘Reactome Pathways 2024’. **c)** Representative immunofluorescence (IF) co-staining for KRT13 (green), TRP63 (red), and KI67 (blue) in SNL squamous tumours and quantification of percent of total Ki67+ cells by co-expression with other marker genes. Representative of n = 4 Ad-CMV-SNL mice. Scale bars, 50 µm. One-way ANOVA. **d)** UMAP of scRNA-seq from lung squamous tumour cells from 18 patients^70^ labeled by Leiden clustering. **e)** UMAP from **4d** labeled by patient (P) sample ID (n = 18). **f-h)** scRNA-seq expression of adenocarcinoma (*NAPSA, NKX2-1, SFTPB*), squamous (*TP63, KRT5, PITX1*), and hillock markers (*KRT13*) in UMAP space from cells in **4d**. **i)** Human hillock score and human basal score (**Supplementary Tables 1 and 2**) applied to tumour cells as in **4d** with score projected in UMAP. **j)** Cell type classification based on hillock score expression from cells in **4d** and KRT13 expression by cell type. Mann-Whitney U test; ****p ≤ 0.0001. **k)** Percent of basal- and hillock-like cells by patient ID as in **4j**. **l)** Gene set enrichment analysis comparing human tumour cells in **4d** and **4j** by assigned cell type. The ENRICHR gene set used was ‘Reactome Pathways 2024’. **m)** CIBERSORTx deconvolution of human LUSC tumours ordered from most to least hillock-like (top); CIBERSORTx deconvolution of leukocyte populations in LUSC tumours ordered same as above (bottom), legend is ordered by correlation with the hillock-like state (see **Methods** and **Supplementary Table 10**). **n)** Cell cycle assignments applied to human tumour cells in UMAP space as in **4d** (see **Methods**), and stacked bar plot showing percent of cells occupying each cell cycle phase by cell state assignment as in **4j.** Statistics shown compare G2/M (orange) or S-phase (green) gene scores between assigned cell states. Mann-Whitney U-Test. ****p ≤ 0.0001. **See also Extended Data Fig. 4 and Supplementary Tables 1-10.**

To determine if a *KRT13^+^* hillock-like state is also present within human squamous tumours, we analyzed single cell data from 18 patient tumours (n = 43,126 cells)^70^ (**Fig. 4d,e**), excluding normal lung cell types such as immune, endothelial, and fibroblast cells (see **Methods**). We analyzed tumour cells for expression of canonical markers of adenocarcinoma (*NAPSA*, *NKX2-1,* and *SFTPB*) and squamous tumours (*KRT5, PITX1* and *TP63*) (**Fig. 4f,g**); as expected, squamous markers were considerably higher than the rarely detectable adenocarcinoma markers. Consistent with observations in SNL tumours, high *KRT13* expression in human tumour cells was associated with an enriched hillock score (**Fig. 4h,i**). While the luminal hillock-score and basal hillock-score largely overlap, the luminal hillock-score more closely mirrors the distribution of *KRT13* expression (**Fig. 4h,i** and **Extended Data Fig. 4e**). Leiden clustering mainly separated cells by patient, likely due to underlying inter-patient genetic and non-genetic heterogeneity, which has been observed in similar studies comparing patient tumours^4,70–73^. However, this limits the ability to identify cells as distinct cell types via Leiden clustering. To circumvent these limitations, we assigned individual cells as the hillock-like state using cutoff criteria defining high hillock-score expression and low basal-score expression (see **Methods**). Using this single-cell classification, hillock-like cells were detected in 17 of 18 human patient samples ranging from ∼1% to 20% of cells per tumour (**Fig. 4j,k** and **Supplementary Table 8**). Similarly, *KRT13*-expressing cells were detected in 16 of 18 human patient samples ranging from ∼2% to 50% of cells per tumour (**Extended Data Fig. 4f** and **Supplementary Table 8**). Together, regardless of classification method, hillock-like cells can be detected at the transcriptional level at frequencies similar to those detected at the protein level in human tissue samples (16 of 20 patient samples) (**Fig. 1e**). To ensure that hillock-like cells share similarities between patients and represent a transcriptionally-unique subset across patient tumours, we performed correlation analysis between basal-like and hillock-like cells from each patient. In most cases, hillock-like tumour cells are more transcriptionally similar to hillock-like cells from other patients than basal-like cells within the same tumour—supporting that the hillock-like state represents a unique transcriptional state conserved across patient tumours (**Extended Data Fig. 4g**).

ENRICHR analysis on DEGs in human hillock-like tumour cells revealed enrichment for immune, neutrophil degranulation, and squamous differentiation signatures including keratinization and cornification (**Fig. 4l** and **Supplementary Table 9**)—similar to the patterns observed in mouse *Krt13*^+^ tumour cells (**Fig. 4a**). We then used the transcriptional profile of the human hillock-like population to perform deconvolution on bulk RNA-seq data of LUSC tumours in TCGA (see **Methods**). Using this approach, we identified a large proportion of LUSCs with cells that resemble the hillock-like population, and many tumours with a high proportion of hillock-like cells (**Fig. 4m**). Deconvolution was used to determine relative proportions of different immune populations in the same tumours. Consistent with our transcriptional predictions and recently published work^65^, tumours with high hillock-like proportions were enriched for immunosuppressive myeloid populations including neutrophils and monocytes, while tumours with a low proportion of hillock-like cells were predicted to have tumour suppressive populations including activated T cells and M1 macrophages (**Fig. 4m** and **Supplementary Table 10**). These data are remarkably consistent with recent spatial transcriptomics of SNL tumours where our colleagues found increased macrophage and monocyte populations in *Tmprss11b*-high hillock-like regions^65^. Cell cycle analysis revealed that human hillock-like tumour cells are relatively less proliferative compared to other cell states in LUSC (**Fig. 4n**), consistent with findings in mice (**Fig. 4c** and **Extended Data Fig. 4b**). Together, these data suggest that a more lowly-dividing KRT13^+^ hillock-like state is present and conserved in both mouse and human LUSC.

### The hillock-like state can arise from NGFR^+^ basal-like tumour cells

To address whether the KRT13^+^ state can arise from the basal-like TPC population, we harvested tumours from Ad-CMV-Cre-infected SNL mice, flow sorted CD45^-^GFP^+^NGFR^+^ tumour cells and plated them in Matrigel to establish organoids (**Fig. 5a**). Importantly, Ad-CMV-derived SNL tumour organoids recapitulate key structures found in SNL squamous tumours *in vivo*, similar to observations by other groups using lung tumour organoid systems^41,43,74–77^. We observed keratinized structures in H&E-stained organoids as well as proliferative KI67^+^ outer edges coinciding with the basal TPC marker, NGFR (**Fig. 5b,c**). Importantly, a KRT13^+^ cell population resided internally within the organoids with limited overlap with the ΔNp63^+^ (also known as P40^+^) basal layer (**Fig. 5d**). Of note, not all organoids had a KRT13^+^ cell population, but all organoids contained basal-like cells, consistent with the emergence of KRT13^+^ cells from basal cells. To confirm the ability of TPC-like cells to regenerate KRT13^+^ populations *in vitro*, we dissociated live tumour organoids and sorted cells into two groups based on surface NGFR expression (NGFR^high^ and NGFR^low^) (**Fig. 5e** and **Extended Data Fig. 5a,** see **Methods**). Sorted cells were replated in Matrigel and allowed to form organoids over 7–10 days in culture. Compared to NGFR^low^ cells, NGFR^high^ cells generated more abundant and larger organoids (**Fig. 5f**). NGFR^low^ cells were capable of forming organoids, suggesting a less basal-like state may possess plasticity to regenerate an NGFR^+^ basal-like state. In order to determine if the NGFR^high^ and NGFR^low^ cells formed organoids with similar structures and cell type distributions, we stained organoids for KRT13 and TP63. Both sorted populations were capable of regenerating organoids containing mutually exclusive TP63^+^ and KRT13^+^ cells (**Fig. 5g**).

**Figure 5.**
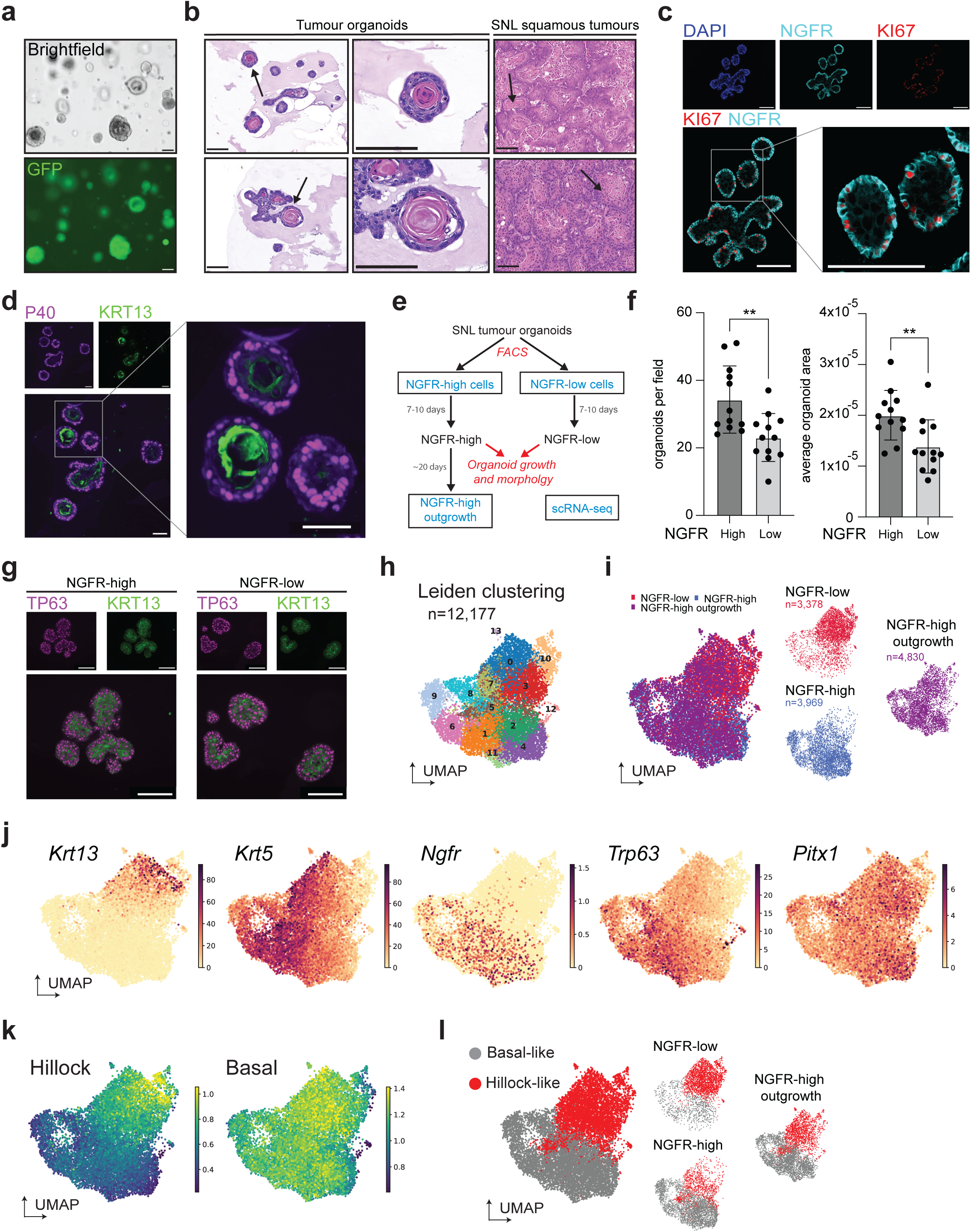
The hillock-like state can arise from NGFR^+^ basal-like tumour cells. **a)** Representative brightfield and immunofluorescence images of tumour organoids established from GFP^+^NGFR^+^ SNL tumour cells. Scale bar, 100 µm. **b)** Representative H&E staining of SNL tumour organoids (n = 2 independent cultures) and SNL squamous tumours (n = 6 mice). Arrows indicate keratinized structures. Scale bars, 100 µm. **c)** Representative immunofluorescence (IF) co-staining for KI67 (red) and NGFR (cyan) in SNL tumour organoids (n = 3). Scale bar, 100 µm. **D)** Representative IF co-staining for P40 (purple) and KRT13 (green) in SNL tumour organoids (n = 3). Scale bar, 50 µm. **e)** Experimental scheme for organoid sorting and scRNA-seq. Populations labeled in blue were used for scRNA-seq. **f)** Number of organoids and average organoid area after SNL tumour organoid cells were sorted by NGFR status and replated in Matrigel and allowed to grow for 10 days. Each point represents an image taken at 4x magnification, and error bars represent mean +/- SD. Two-tailed t-test, **p ≤ 0.01. **g**) Representative IF co-staining for TP63 (purple) and KRT13 (green) from SNL tumour organoids allowed to grow for 10 days after sorting. Representative of n = 2 experiments with 6 different 20x fields per experiment. **h)** Leiden clustering of scRNA-seq performed on SNL tumour organoids as in **5e**. **i)** scRNA-seq of SNL tumour organoids at time of NGFR sorting (NGFR-high and NGFR-low) and NGFR-high cells after four weeks of growth in organoid conditions (“NGFR-high outgrowth”) in **5h**. **j)** Expression of indicated hillock (*Krt13*) or basal (*Ngfr*, *Krt5*, *Trp63, Pitx1*) genes in UMAP space from cells in **5h**. **k)** scRNA-seq expression of the hillock score and basal score projected on UMAP space from cells in **5h. l)** Cell type classification by Leiden clustering based on gene and gene signature score expression from cells in **5h**. (see **Methods** and **Supplementary Table 1 and 2**). **See also Extended Data Fig. 5**.

To determine the transcriptional heterogeneity of tumour organoids as well as the ability of basal-like cells to give rise to hillock-like cells, we performed scRNA-seq on freshly dissociated NGFR^high^ and NGFR^low^ organoid cells as well as sorted NGFR^high^ cells that were allowed to regrow as organoids for four weeks in culture (“NGFR^high^ outgrowth”) (**Fig. 5e** and **Extended Data Fig. 5a**). Importantly, organoids largely transcriptionally recapitulate the heterogeneity observed in squamous SNL tumours, with distinct populations of *Ngfr/Trp63*-high basal-like cells and *Krt13/Tmprss11b*-high hillock-like cells (**Fig. 5h-j** and **Extended Data Fig. 5b**). Consistently, there was a subpopulation of hillock-score-enriched cells within organoids, while the basal score was highly expressed across the entire transcriptional space (**Fig. 5k**). The abundance of basal-like states in organoid conditions is not surprising, as other studies have shown that the Matrigel organoid culture conditions support basal-like states^69,78^. Basal hillock and luminal hillock signatures overlapped with the general basal and hillock signatures, respectively (**Fig. 5k** and **Extended Data Fig. 5c**), similar to patterns in SNL tumour data (**Fig. 3g**). Given that mucinous histopathology precedes squamous differentiation in vivo^18,30,40,49–51^, we did not expect to observe mucinous-like cells in organoids, and consistently, mucinous markers were extremely rare and low (**Extended Data Fig. 5c**). Outgrowth of NGFR^high^ cells repopulated the entirety of transcriptional space occupied by both freshly-sorted NGFR^high^ and NGFR^low^ cells, suggesting that basal cells can regenerate all captured cell states (**Fig. 5i**). Additionally, *Krt13* expression and the hillock score were restricted to NGFR^low^ (and not NGFR^high^) cells at the time of sorting (**Extended Data Fig. 5d**). Outgrowth of NGFR^high^ cells led to reappearance of *Krt13* expression and a hillock-like population (**Fig. 5l** and **Extended Data Fig. 5d**), together supporting that hillock-like cells can derive from basal-like TPCs. Consistent with immunostaining for KI67 (**Fig. 5c**), hillock-like organoid cells are less proliferative based on cell-cycle gene expression assignment (**Extended Data Fig. 5e**). Organoid hillock-like cells also upregulate neutrophil degranulation and cornification gene sets (**Extended Data Fig. 5f** and **Supplementary Table 11**), similar to both mouse and human tumour data.

### KLF4 is enriched in a suprabasal hillock-like state in squamous tumours

Next, we sought to identify master transcriptional regulators driving the hillock-like tumour state. We assessed the ENCODE and ChEA transcription factor gene sets in ENRICHR using highly expressed genes across our mouse, human, and organoid single-cell datasets (**Extended Data Fig. 6a-c** and **Supplementary Table 12**). Basal- and basal hillock-like cells across mouse and human datasets predicted proliferation and basal-lineage transcription factors MYC, MAX, P63 and SOX2 (**Extended Data Fig. 6a**), consistent with their increased proliferation compared to luminal hillock-like cells. In contrast, KLF4 was a top predicted regulator of both mouse and human hillock-like tumour cells and was also a top predicted regulator of normal airway hillock cells from two independent datasets^6,7^ (**Extended Data Fig. 6b, c**). KLF4 is a promising candidate for regulating the hillock-like state because of its role in multiple squamous epithelial tissues (i.e., esophagus, skin) in promoting cell differentiation away from the basal lamina^55,79^. Moreover, KLF4 can directly regulate *KRT13* expression in esophageal cancer^80^ and can repress basal TPC factors SOX2, TP63 and PITX1^23^. To determine KLF4’s expression pattern in lung squamous tumours, we queried our RNA-seq data from lung cancer GEMMs and TCGA data, which showed enrichment of *Klf4/KLF4* in both mouse and human squamous tumours compared to adenocarcinomas (**Fig. 6a,b**).

**Figure 6.**
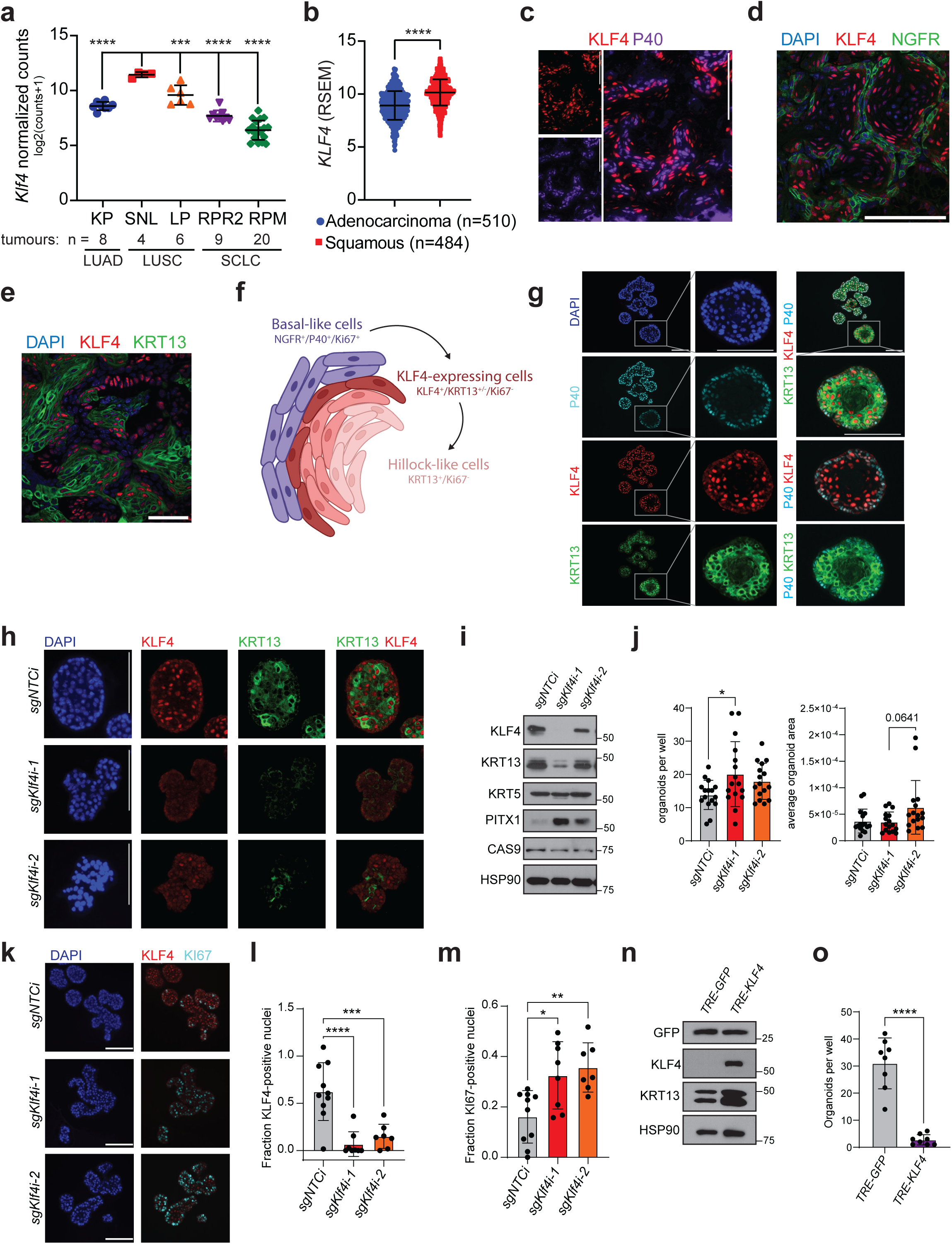
KLF4 is enriched in a suprabasal hillock-like state in squamous tumours and regulates KRT13 expression. **a)** Log2 normalized counts of *Klf4* from RNA-seq of lung tumours from indicated GEMMs with number of tumours indicated in the figure. Statistical significance determined by one-way ANOVA. **b)** RSEM expression of *KLF4* from bulk human RNA-seq data of lung adenocarcinoma and squamous cell carcinoma from TCGA with number of samples indicated in the figure. Two-tailed t-test. **c)** Representative immunofluorescence (IF) co-staining for KLF4 (red) and dNP63/P40 (purple) in squamous tumours from Ad-CMV-initiated SNL tumours. Representative of n = 2 mice. Scale bar, 100 µm. **d)** IF staining for NGFR (green) and KLF4 (red) in squamous tumours from Ad-CMV-initiated SNL tumours. Representative of n= 2 mice. Scale bars, 100 µm. **e)** Representative IF co-staining for KRT13 (green) and KLF4 (red) in squamous tumours from Ad-CMV-initiated SNL tumours. Representative of n= 3 mice. Scale bars, 50 µm. **f)** Schematic of a squamous tumour with basal-like cells, KLF4-expressing suprabasal cells, and KRT13-expressing hillock-like cells. Made with Biorender. **g)** Representative IF co-staining for KRT13 (green), KLF4 (red), and dNP63/P40 (cyan) in SNL tumour organoids. Scale bars, 100 µm. Representative of n = 3 experiments with at least 5 different 20x fields per experiment. **h)** Representative IF co-staining for KRT13 (green) and KLF4 (red) in SNL tumour organoids with KLF4 knockdown using CRISPRi. Scale bars, 100 µm. Representative of n = 2 experiments at least 4 different 20x fields per experiment. **i)** Immunoblot for indicated proteins from SNL tumour organoids after 10 days of growth following dissociation to single cells and replating in Matrigel. **j)** Number and average size of organoids after 7 days of growth following dissociation to single cells and replating in Matrigel. One-way ANOVA. **k)** Representative IF co-staining for KI67 (cyan) and KLF4 (red) in SNL tumour organoids with KLF4 knockdown using CRISPRi. Scale bars, 100 µm. Representative of n = 2 experiment with at least 7 different 20x fields per sample. **l-m)** Quantification of nuclei staining positive for KLF4 (**l**) or KI67 (**m**) from **6k**. One-way ANOVA. **n)** Immunoblot for indicated proteins from SNL organoids using the TRE-GFP or TRE-KLF4 vectors. Cells were grown for 10 days and 1µg/mL doxycycline was added every 48 hrs. **o)** Number and average size of SNL tumour organoids after 7 days of growth following dissociation to single cells and replating in Matrigel and treated with doxycycline as in **6n**. Two-tailed t-test. **For all panels**, each data point represents an individual tumour or a 4x (organoid growth)/20x (IF quantification) microscopy field of view and error bars represent mean +/- SD; *p ≤ 0.05;**p ≤ 0.01; ***p ≤ 0.001; ****p ≤ 0.0001. DAPI (blue) labels nuclei. HSP90 serves as loading control for immunoblots. **See also Extended Data Fig. 6.**

To determine KLF4 expression and localization in LUSC, we co-stained for basal marker ΔNp63/P40 and KLF4 in mouse SNL tumours. KLF4 exhibited a nuclear and suprabasal-like pattern of expression where KLF4 was closely positioned to, but largely non-overlapping with P40^+^ cells (**Fig. 6c**). KLF4 was similarly located in a suprabasal pattern when co-staining with the basal TPC-marker NGFR in both Ad-CMV- and Ad-CCSP-initiated tumours (**Fig. 6d** and **Extended Data Fig. 6d**). Co-staining for KLF4 and KRT13 showed that KLF4 was highly expressed in a layer of KRT13^+^ cells (typically in the outermost KRT13^+^ basal-proximal cells), and absent from the innermost KRT13^+^ basal-distal cells (**Fig. 6e,f** and **Extended Data Fig. 6d**). Finally, we stained SNL organoids for KLF4 and noted KLF4 expression enriched in a suprabasal location to NGFR^+^/P40^+^ cells, with minimal overlap between KLF4 and basal markers (**Fig. 6g** and **Extended Data Fig. 6e**); KLF4 was expressed in the majority of KRT13^+^ cells, suggesting that organoids have a similar organization as squamous tumours. These data suggest multiple levels of intra-tumoural heterogeneity in squamous lung tumours (**Fig. 6f**) that mimic normal stratified epithelia, similar to findings in the skin and vaginal epithelium^57,81^. These data suggest KLF4 is positioned to be important for the emergence or differentiation of KRT13^+^ hillock-like lung tumour cells.

### KLF4 promotes a KRT13^+^ hillock-like state

To determine the functional role of KLF4 in the hillock-like tumour state, we sought to deplete KLF4 expression and determine the impact on KRT13 and the hillock-like state in SNL tumour organoids. Multiple attempts to knockout *Klf4* using traditional CRISPR gene-editing were unsuccessful due to organoid death. Thus, we designed a CRISPR-inhibition (CRISPRi) system using a catalytically dead Cas9 enzyme (dCas9) fused to the transcriptional repressor KRAB protein^82^. Compared to non-targeting control sgRNAs (sgNTCi), *Klf4*-targeted CRISPRi constructs (sg*Klf4*i) efficiently decreased KLF4 expression in SNL organoids, especially guide #1 (**Fig. 6h,i**). Analysis of sgNTCi and sg*Klf4*i organoids by co-IF and immunoblot revealed significantly reduced and more diffuse KRT13 staining upon *Klf4* knockdown (**Fig. 6h,i**). The CRISPRi system led to variable KLF4 expression within organoids of the same culture, where KRT13 expression was retained in KLF4-proficient sg*Klf4i* organoids but completely absent in KLF4-deficient sg*Klf4i* organoids (**Extended Data Fig. 6f**). KLF4 suppression did not alter expression of the basal marker KRT5 but increased expression of PITX1 (**Fig. 6i**), a transcription factor that cooperates with SOX2 and TP63 in basal skin TPCs, with a mutually antagonistic relationship with KLF4^23^. After seven days of growth following dissociation and plating, KLF4 knockdown with sg*Klf4i-1* slightly but significantly increased organoid frequency with no impact on organoid size (**Fig. 6j**). Accordingly, sg*Klf4i* organoids showed elevated KI67 staining, with KI67^+^ cells present throughout the organoid when KLF4 was suppressed, whereas KLF4-proficient organoids tended to proliferate specifically in the basal-associated outer layer of cells (**Fig. 6k-m**). These data may be consistent with known functions of KLF4 in regulating cell cycle exit^23,54,83–85^. The lack of increased organoid size despite increased proliferative markers may reflect size constraints in culture conditions. Together, these data suggest that KLF4 is necessary for the transition to a less proliferative KRT13^+^ hillock-like state in the organoid context.

Next, we aimed to determine whether KLF4 is sufficient to promote the KRT13^+^ hillock-like state. We overexpressed human *KLF4* or *GFP* control in SNL organoids under the control of a doxycycline-inducible tetracycline response element (Tet-ON-TRE-GFP vs KLF4). KLF4 overexpression in SNL tumour organoids increased KRT13 levels and substantially reduced organoid growth (**Fig. 6n,o**). After dissociation and re-plating with doxycycline for 7 days, organoids with KLF4 overexpression were smaller and formed at a lower frequency than GFP controls (**Fig. 6o** and **Extended Data Fig. 6g**). Together, these data suggest KLF4 is both necessary and sufficient for the emergence of the KRT13^+^ state in SNL organoids.

We extended these findings to human squamous lung cancer-derived cell lines that express varying degrees of KLF4 and KRT13 at baseline (**Extended Data Fig. 7a**). Consistent with findings in murine organoids, *KLF4* knockdown in human LUSC led to a decrease in KRT13 (**Fig. 7a** and **Extended Data Fig. 7b**), while overexpression of KLF4 induced KRT13 (**Fig. 7b**). As in SNL organoids, KLF4 overexpression suppressed growth of H226 cells in vitro (**Fig. 7c**). In contrast to tumour organoids where KLF4 loss increased proliferation (**Fig. 6i-m**), KLF4 knockout in H520 cells decreased proliferation (**Extended Data Fig. 7c**), which may be due to unique constraints in these distinct cell populations, which differ by species, genetic alterations, and in vitro growth conditions.

**Figure 7.**
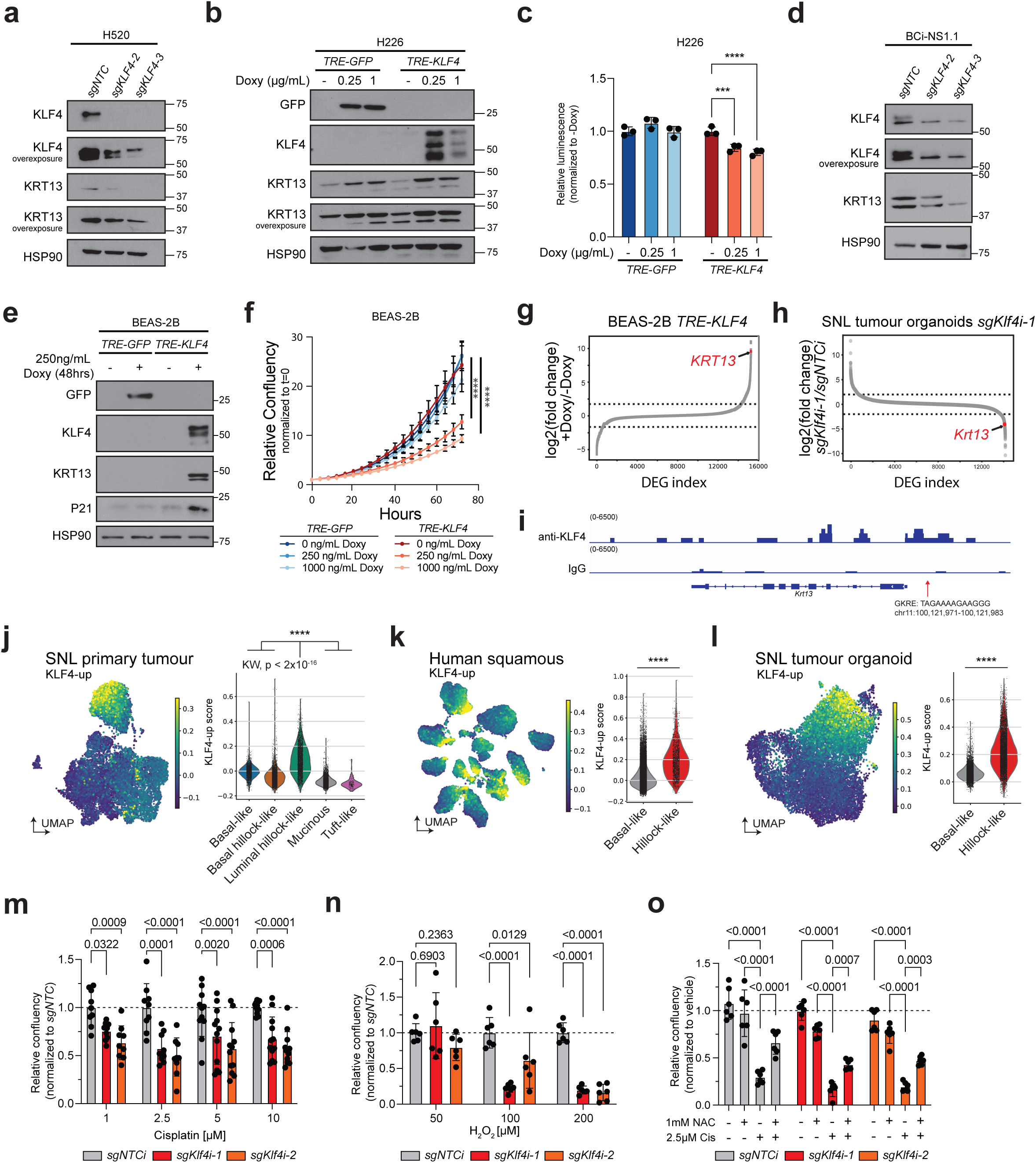
KLF4 promotes a KRT13^+^ hillock-like state and resistance to cisplatin. **a)** Immunoblot for indicated proteins in the human LUSC cell line H520 expressing non-targeting control (*sgNTC*) or *sgKLF4*. **b)** Immunoblot for indicated proteins in the human H226 LUSC cells with *TRE-GFP* or *-KLF4* overexpression constructs at the indicated dose or without (-) doxycycline (doxy) exposure for 48 hrs. **c)** Celltiter-Glo (Promega) luminescence normalized to no doxy (-) after 5 days of growth in H226 cells treated with the indicated dose of doxy every 48 hrs. Two-way ANOVA, comparing within genotype only. **d)** Immunoblot for indicated proteins in human BCi-NS1.1 cell lines expressing non-targeting control (*sgNTC*) or *sgKLF4*. **e)** Immunoblot for indicated proteins in human BEAS-2B cell lines with *TRE-GFP* or *-KLF4* overexpression constructs with (+) or without (-) doxycycline exposure for 48 hrs. **f)** Cell growth determined by confluency using an Incucyte live-cell imaging system (Sartorius) normalized to initial plating density of BEAS-2B cells grown with varying doses of doxycycline (Doxy). One-way ANOVA. **g)** Waterfall plot of differentially expressed genes from RNA-sequencing of TRE-KLF4 BEAS-2B cells treated +/-doxy as in **7f**. **h)** Waterfall plot of differentially expressed genes from RNA-sequencing of SNL tumour organoids comparing *sgKlf4i-1* to *sgNTC* from **6h-i**. **i)** IGV genome viewer of the *Krt13* locus for KLF4 CUT&RUN reads in SNL tumour organoids. **j-l)** KLF4-up score applied to SNL tumour cells (**j**), human squamous tumour cells (**k**), and organoid cells (**l**) projected onto UMAP space from **3a, 4d,** and **5h** with score expression violin plots by cell state assignment. Kruskal-Wallis followed by Dunn’s test with a Bonferonni correction; all comparisons p < 2×10^-16^ (**j**), Mann-Whitney U test (**k,l**); ****p ≤ 0.0001. (see **Methods** and **Supplementary Table 13**). **m)** Organoid growth after treatment with cisplatin determined by GFP+ confluency using an Incucyte live-cell imaging system (Sartorius) normalized to vehicle treated organoids followed by normalization to *sgNTCi* after 5 days. Two-way ANOVA, multiple comparisons to *sgNTCi* within each dose. **n)** Organoid growth after treatment with hydrogen peroxide (H_2_O_2_) determined by GFP+ confluency using an Incucyte live-cell imaging system (Sartorius) normalized to vehicle treated organoids followed by normalization to *sgNTCi* after 5 days. Two-way ANOVA, multiple comparisons to *sgNTCi* within each dose. **o)** Organoid growth after treatment with cisplatin and N-acetylcystine (NAC) determined by GFP+ confluency using an Incucyte live-cell imaging system (Sartorius) normalized to vehicle treated organoids after 5 days. Vehicle treatment represented by dashed line at relative growth 1 for each cell line. Two-way ANOVA, multiple comparisons to vehicle within each genotype. **For all panels,** error bars represent mean +/- SD; *p ≤ 0.05; **p ≤ 0.01;***p ≤ 0.001; ****p ≤ 0.0001 or exact values are shown. DAPI (blue) marks nuclei. HSP90 serves as loading control for immunoblot. **See also Extended Data Fig. 7 and 8.**

Given that basal cells are implicated as the cell of origin for LUSC^32^, that basal hillock cells give rise to luminal hillock cells in the lung airways^37^, and that basal TPC-like cells differentiate into hillock-like tumour cells, we asked if KLF4 can modulate KRT13 in normal lung basal cells. We used CRISPR-Cas9 gene editing to knockout *KLF4* in BCi-NS1.1 cells, which naturally express KLF4 and have been shown to undergo differentiation away from a basal state in cell culture conditions^86–88^. (**Fig. 7d** and **Extended Data Fig. 7d**). As in tumour organoids and H520 human LUSC cells, *KLF4* knockout substantially decreased endogenous KRT13 levels especially with guide #3, which showed the most complete DNA editing by surveyor assay (**Fig. 7d** and **Extended Data Fig. 7d**). Similar to findings in H520 cells, *KLF4* knockout in BCi-NS1.1 cells led to reduced proliferation (**Extended Data Fig. 7e**), again suggesting context specific functions in 2D cell culture compared to 3D organoid culture.

To determine if KLF4 is sufficient to promote a KRT13^+^ state in human cells, the TRE-KLF4 overexpression construct was introduced into BEAS-2B cells that are considered a bronchial epithelial cell type that lacks expression of KLF4 and KRT13 under normal culture conditions (**Fig. 7e** and **Extended Data Fig. 7f**). Exposure to doxycycline induced KLF4 and led to striking KRT13 induction (**Fig. 7e**). Additionally, transient exposure of TRE-KLF4 cells to doxycycline was able to induce KRT13 expression, which was maintained for 48 hrs after doxycycline was removed, while KLF4 expression returned to baseline after doxycycline removal (**Extended Data Fig. 7f**). KLF4 overexpression significantly decreased growth of BEAS-2B cells (**Fig. 7f**), which is consistent with the slower proliferating KRT13^+^ hillock-like state observed in LUSC tumours and organoids (**Fig. 4c, 5c, 6k-o** and **7c**). KLF4 overexpression also led to increased P21 (**Fig. 7e**), consistent with previous work^54,83–85^.

Transcriptomic analysis of BEAS-2B cells following KLF4 induction revealed significant upregulation of genes associated with the hillock state, including *KRT13* and genes associated with *Krt4/Krt13+* mouse tracheal cells, and cornification and keratinization-related genes (**Fig. 7g, Extended Data Fig. 7g,h** and **Supplementary Table 13**). Similarly, transcriptomic analysis of SNL tumour organoids with KLF4 silenced (sg*Klf4i*-1) showed suppression of *Krt13* and hillock-associated genes (**Fig. 7h, Extended Data Fig. 7i,j,** and **Supplementary Table 13**). Notably, KLF4 overexpression and knockdown did not appear to alter immune-related gene sets, suggesting that KLF4 may regulate a part of the hillock-like program including *Krt13/KRT13* but not all hillock-associated genes (**Extended Data Fig. 7g-j**). To determine if KLF4 induction of *Krt13* and other hillock-associated genes is through direct transcriptional regulation, we performed KLF4 CUT&RUN in SNL tumour organoids. Annotated peaks were enriched for the KLF4 binding motif as expected (**Extended Data Fig. 7k**). A KLF4 bound region was detected directly upstream of the *Krt13* locus (**Fig. 7i**), consistent with a previously annotated GKLF/KLF4-responsive element (GKRE)^80^. Additionally, ENRICHR analysis of the top 500 promoter regions of protein-coding genes bound by KLF4 showed enrichment of genes involved in Rho GTPase signaling (known to regulate squamous epithelial differentiation), neutrophil degranulation, and Ephrin signaling, signatures seen in our single cell analysis across models. The bound promoters were also enriched for genes associated with normal *Krt13+* hillock cells of the mouse trachea (**Extended Data Fig. 7l-m** and **Supplementary Table 14**).

Finally, we generated a conserved 51-gene score between the overlapping genes upregulated in BEAS-2B cells following KLF4 induction and downregulated by *Klf4* knockdown in SNL tumour organoids (“KLF4-up score”) and applied this score to our mouse and human single cell datasets (**Extended Data Fig. 7n** and **Supplementary Table 13**, see **Methods**). Consistently, KLF4-induced genes were highly enriched in the hillock-like populations in all three single cell datasets (**Fig. 7j-l**), supporting a role for KLF4 in hillock-like cancer cells. Together, these data suggest that KLF4 promotes KRT13 and a hillock-like state in both normal lung cell lines and tumour contexts.

### KLF4 promotes resistance to chemotherapy

Hillock cells of the normal airways are highly resistant to multiple forms of injury including viral infection and physical stress^37^. We hypothesized that hillock-like cells in squamous tumours may possess similar injury-resistant phenotypes. Focusing on standard-of-care chemotherapies cisplatin and paclitaxel, we assessed if the hillock-like state dictates response to clinically-relevant treatments. Organoid cells with and without KLF4 knockdown were treated with varying doses of either cisplatin or paclitaxel as single-cell suspensions, replated with fresh drug, and allowed to form organoids. Cisplatin decreased organoid proliferation within five days, with *sgKlf4i-*organoids displaying increased sensitivity versus *sgNTCi* controls at all doses tested (**Fig. 7m**). Additionally, while control organoids were able to repopulate during extended culture following 5 µM cisplatin, *sgKlf4i* organoids were not (**Extended Data Fig. 8a**). At a lower dose of 2.5 µM cisplatin, *sgKLF4i-1* organoids repopulated but exhibited restored expression of KLF4 and KRT13, suggesting escape from KLF4-knockdown may facilitate cisplatin resistance (**Extended Data Fig. 8a,b**). Interestingly, KLF4 knockdown had no effect on organoid response to paclitaxel (**Extended Data Fig. 8c**). Similar to findings in mouse organoids, KLF4-knockout increased sensitivity to cisplatin in the human basal cell line BCi-NS1.1, while having no effect on paclitaxel response (**Extended Data Fig. 8d-f**). BCi-NS1.1 cells with the most complete KLF4-knockout (*sgKLF4-3,* **Fig. 7d**) exhibited the most profound sensitivity to cisplatin with evidence of apoptosis as measured by cleaved PARP and cleaved Caspase-3 levels (**Extended Data Fig. 8e**).

Of note, a top predicted regulator of our hillock-like state is *NFE2L2* (NRF2) (**Extended Data Fig. 6b, c**), a master regulator of oxidative stress. Cisplatin is known to induce oxidative stress along with DNA damage, whereas this is not a well annotated response to paclitaxel. Thus, we hypothesize that selective sensitization to cisplatin and not paclitaxel may be partially mediated by oxidative stress. Indeed, organoids harbouring KLF4 loss were dramatically sensitized to treatment with hydrogen-peroxide-induced oxidative stress compared to controls (**Fig. 7n**). Further, treatment of cells with the antioxidant N-acetylcysteine (NAC) provided a partial rescue of organoid proliferation in response to cisplatin (in both control and knockdown settings) (**Fig. 7o**). Recent work has suggested NRF2 and KLF4 cooperate to regulate oxidative stress response genes^89^, and NRF2 has been implicated in resistance to numerous therapies^41,90,91^, such that future studies should address whether NRF2 cooperates with KLF4 in the hillock-like state. Together, these data suggest that KLF4 supports resistance to chemotherapy, associated with its capacity to drive a hillock-like state.

### A conserved hillock-like state is present across squamous tumours with potential therapeutic targets

Squamous cell carcinomas (SCCs) can arise in tissues beyond the lung and display similar histologic features^92–94^. Additionally, there are common underlying genetic alterations that promote squamous tumours, especially in the lung, head & neck, and esophageal contexts^92–94^. We therefore queried two previously published SCC datasets, one containing 12,932 cells from 15 patients with head & neck SCC (HNSCC)^95^ and one containing 44,547 cells from 60 patients with esophageal SCC (ESCC)^96^ (**Fig. 8a** and **Extended Data Fig. 9a**), for evidence of the hillock-like state. Both tumour types contained populations of cells that expressed *KRT13*, as well as the basal markers *KRT5* and *TP63*, with minimal overlap between them (**Extended Data Fig. 9b**). The basal score was broadly expressed across the datasets while the hillock score was more restricted to clusters that are lower for basal score expression (**Extended Data Fig. 9c**), similar to patterns seen in LUSC (**Fig. 4i**). Using methods identical to those used for the human LUSC analyses, we assigned cell states to both the HNSCC and ESCC cells using a hillock score cutoff approach (**Fig. 8b** and see **Methods**). Hillock-like cells in both datasets showed evidence of decreased proliferation by cell cycle phase assignment; ENRICHR analysis of upregulated genes in the hillock-like state showed remarkable similarity to LUSC, including innate immune and squamous-related gene signatures (**Extended Data Fig. 9d,e**). Finally, the KLF4-up score was enriched in hillock-like cells in both HNSCC and ESCC (**Fig. 8c**). Together, these findings suggest that a KLF4-driven hillock-like state exists across multiple squamous tumour types and that understanding this cell state should have implications beyond lung cancer.

**Figure 8.**
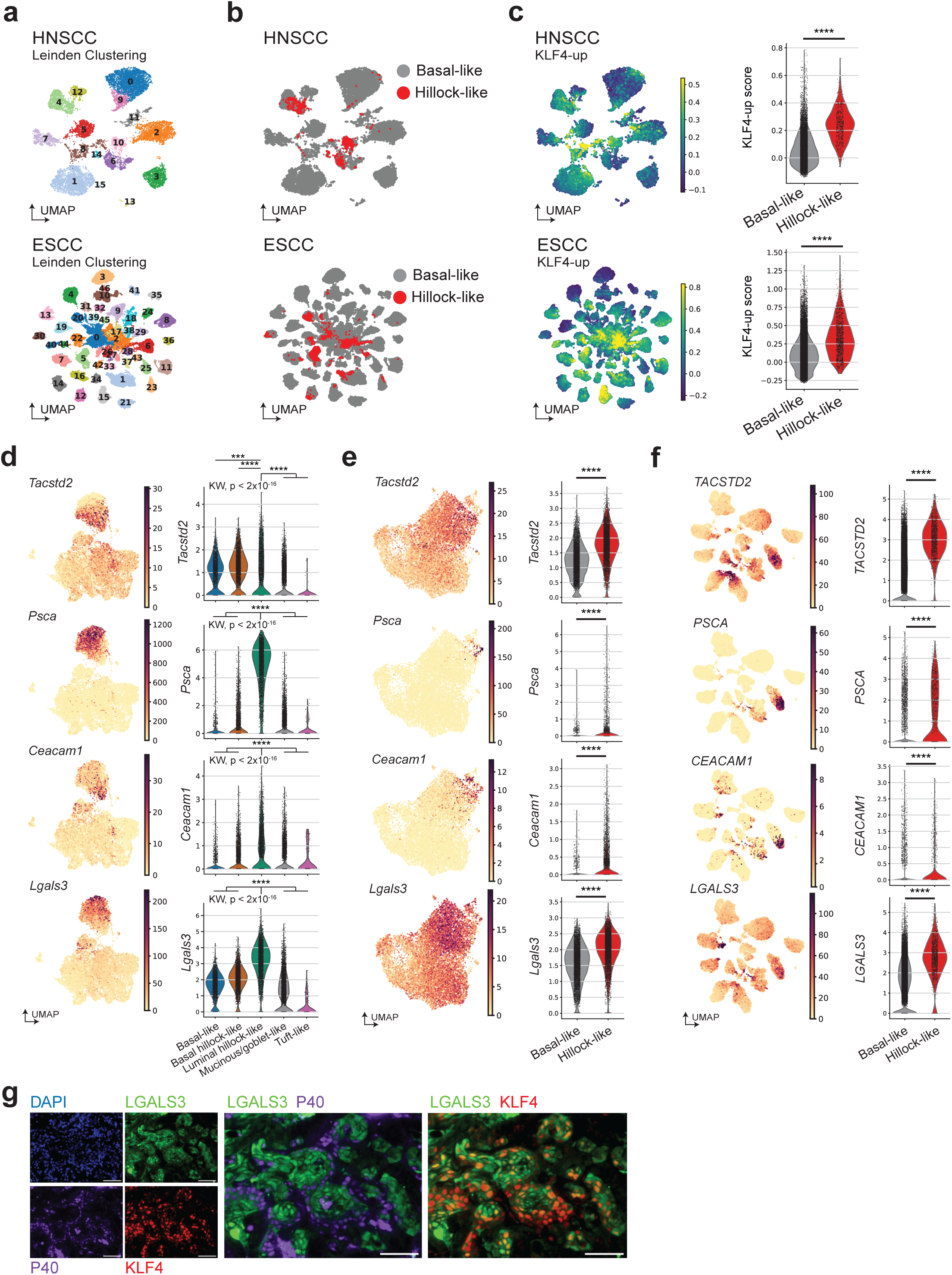
A conserved hillock-like state is present across squamous tumours and is enriched for potential therapeutic targets. **a)** UMAP of scRNA-seq from head & neck tumour cells from 15 patients^95^ (Kurten et al) or from esophageal tumours from 60 patients^96^ (Zhang et al) labeled by Leiden clustering. **b)** Cell type classification of head & neck SCC or esophageal SCC single cell data based on hillock score expression from cells in **8a**. **c)** UMAP of KLF4-up score applied to head & neck and esophageal SCC cells from **8a** with score expression in violin plots grouped by cell state assignment. Mann-Whitney U test; ****p ≤ 0.0001. **d-f)** Gene expression in SNL tumour cells (**d**), organoid cells (**e**), and human squamous tumour cells (**f**) projected onto UMAP space from **3a, 4d,** and **5h** with score expression violin plots by cell state assignment. Kruskal-Wallis followed by Dunn’s test with a Bonferonni correction; all comparisons p < 2×10^-16^ (**d**), Mann-Whitney U test (**e,f**); ****p ≤ 0.0001. **g)** Representative immunofluorescence (IF) co-staining for LGALS3 (green), KLF4 (red), and P40 (purple) in SNL squamous tumours. Representative of n = 2 Ad-CMV-SNL mice. Scale bars, 50 µm. **See also Extended Data Fig. 9 and 10.**

The ability to differentiate and target specific tumour cell states could allow the ability to study the functional role of individual cell states in disease progression. KRT13 appears to be a marker of the hillock-like state across models and tissues, which may have utility for potential diagnostic purposes. To identify additional clinically-relevant markers, we sought to identify hillock-enriched cell surface markers, which could act as potential targets for antibody drug conjugates (ADCs) or bi-specific T cell engagers (BiTes). We identified *TACSTD2* (encoding TROP2) as a top cell surface protein-encoding gene enriched in hillock-like cells across models (**Fig. 8 d-f**; **Extended Fig. 10a-d** and **Supplementary Table 15**) as well as *PSCA* and *CEACAM1*. TROP2 ADCs have recently been approved for treatment of breast cancer and EGFR-mutated lung cancer and are in numerous clinical trials (NCT02574455, NCT03901339, NCT07328490, TROPION-Lung, etc). Finally, *Lgals3* is a top hillock-enriched gene throughout our study (**Extended Data Fig. 4a, Supplementary Table 6** and **7**), and while not a cell-surface protein per se, it is a secreted lectin that localizes to the cell surface via interactions with glycosylated receptors (**Fig 8d-f**). We validated LGALS3 to be restricted to the hillock-like compartment in SNL tumours by IF (**Fig. 8g**). LGALS3 has been implicated in neutrophil recruitment and regulation of pro-tumor immunity^97–101^, while drugs targeting it have good safety profiles in early-stage clinical trials^102^. Future studies are warranted to determine if these targets can be exploited to enrich for the hillock-like state, evaluate its presence in patient samples, and/or develop novel therapies specifically targeting hillock-like cells.

## Discussion

Here, we identify a KRT13⁺ hillock-like tumour cell state in LUSC that resembles recently described hillock states in the normal airway epithelium^37^. Hillock-like tumour cells are present across mouse tumours, tumour organoids, and human LUSC, and we detect similar populations in additional squamous malignancies. Together, our findings support the idea that squamous tumours exhibit lineage plasticity, adopting transcriptional states that mirror normal epithelial cell states, and that these programs contribute to the non-genetic heterogeneity of LUSC.

Our data support a model in which hillock-like cells represent a basal-derived state that is more differentiated and slower dividing. In the normal airway, hillock cells are linked to epithelial protection and regeneration after injury^6,37^; in tumours, a similar program may allow a subset of cells to persist under stress while maintaining the capacity to transition between states. This raises key questions for future studies, including whether basal-like cells transition into hillock-like states in vivo, whether hillock-like cells can revert back to basal-like populations after therapy, and which microenvironmental signals promote or restrict these transitions.

We also find that cell of origin influences hillock-like programs in the SNL model. Club-initiated tumours more prominently exhibit luminal hillock-like features, whereas basal-enriched contexts (Ad-CMV-Cre and tumour organoids) more readily show basal-hillock-like features, reminiscent of lineage-tracing studies showing that normal hillock cells are intermediate between basal and club cells^6,7^. Together, these findings suggest that lineage history and epigenetic context may shape which cell states are most accessible during tumour evolution. This paradigm, in which the cell of origin impacts lineage trajectories, has also been observed in small cell lung cancer^69^ and is therefore likely broadly relevant across cancers.

Mechanistically, our data identify KLF4 as a key regulator of the hillock-like state. KLF4 directly binds the *Krt13* locus and regulates a conserved KLF4-driven transcriptional module enriched in hillock-like populations across mouse and human contexts. At the same time, hillock-like cells display broader immunomodulatory features that may not be fully explained by KLF4 alone, suggesting that additional regulators contribute to this program. Another top candidate predicted to regulate the hillock-like state from our computational analyses is NRF2, which is known to regulate oxidative stress responses, the tumour immune microenvironment, and therapeutic resistance in LUSC, all phenotypes of hillock-like cancer cells based on our findings^41,90,91,103^. Future work should test whether KLF4 cooperates with NRF2 or other factors to shape the hillock-like phenotype, including immune-related features.

The potential clinical relevance of this state is supported by the finding that KLF4 promotes resistance to cisplatin and oxidative stress. A slower-dividing, stress-resistant cell population could survive cytotoxic therapy and contribute to tumour persistence and regrowth through plasticity. In addition, hillock-like programs correlate with immune contexts enriched for myeloid populations in human datasets, which we and colleagues also observe in SNL lung tumours using spatial transcriptomics^65^. TMPRS11B, which is expressed in *Krt13*+ hillock-like cells and is regulated by KLF4^65^, drives lactate export by promoting MCT4 activity and glycolytic metabolism, which correlates with low-pH regions and increased innate immune infiltration, providing one example of how the hillock-like state may alter the tumour immune microenvironment^64,65^. Together, these findings are consistent with the possibility that specific tumour cell states shape the immunosuppressive microenvironment in distinct ways. Future studies should directly test whether hillock-like tumour cells influence immune recruitment or function, and whether these interactions impact response to immunotherapy.

Finally, our analyses highlight potential markers and vulnerabilities enriched in hillock-like cells, including therapeutically tractable surface-associated targets such as TACSTD2 (TROP2), CEACAM1, PSCA, and LGALS3. Together, these observations motivate future work to develop strategies that selectively track, enrich, or target hillock-like tumour cells. Overall, our findings suggest that effective LUSC therapy may require targeting both proliferative basal-like populations and stress-adapted states that support tumour plasticity, therapy resistance, and immune suppression.

### Limitations of the Study

Our study suggests non-genetic factors support intra-tumoural heterogeneity in LUSC and that *Krt13^+^* hillock-like cells may contribute to tumour progression, but we acknowledge limitations to our systems. First, our mouse models and organoids are predominantly derived from the SNL model. *SOX2* amplification is one of the most common genetic events in squamous tumours and SNL squamous tumours are known to recapitulate many features of human LUSC^40^; however, this model harbours mucinous adenocarcinomas, and squamous tumours are likely derived from transdifferentiation from an adenocarcinoma state, which may not reflect the origins of many human tumours. Because we are not yet able to resolve a luminal-hillock-like and basal-hillock-like cell state in human tumour data (partially due to higher levels of heterogeneity limiting clustering-based approaches), we cannot be sure whether human LUSC is predominantly basal and basal-hillock, or contains luminal hillock-like populations as well.

Reliance on one GEMM limits the ability to assess the impact of genetics on the hillock-like state. We show that the LP model and human tumours also contain KRT13-expressing cells with transcriptional similarities between other human squamous tumours, but future work should incorporate additional genetic models. Our findings suggest that cell of origin impacts intra-tumoural heterogeneity, as tumours initiated in club cells show increased KRT13 positivity compared to Ad-CMV-Cre-initiated tumours. Given that the cells of origin with Ad-CMV-Cre are unclear, further characterization using more specific Cre drivers (including a basal *Krt5*-Cre) can clarify these developmental relationships.

## AUTHOR CONTRIBUTIONS

Conceptualization: L.T. Izzo, K.A. O’Donnell, T.G. Oliver

Data Acquisition: L.T. Izzo, T. Reyes, S. Meesala, S. Yang, H.S. Sunil, A.S. Ireland, L. Earnest-Nobel, X.C. Cheng, N. Tserentsoodol, S.B. Hawgood

Data Analysis and Interpretation: L.T. Izzo, T. Reyes, H.S. Sunil, A.S. Ireland, C. Glass, D.R. Tyson, T.G. Oliver

Writing – Original Draft: T. Reyes, L.T. Izzo, T.G. Oliver

Writing – Review and Editing: L.T. Izzo, S. Meesala, A.S. Ireland, T. Reyes, D.R. Tyson, K.A. O’Donnell, T.G. Oliver

Funding Acquisition: L.T. Izzo, K.A. O’Donnell, T.G. Oliver

Resources: E.F. Patz, B.J. Witt

Supervision: K.A. O’Donnell, T.G. Oliver

## ACKNOWLEDGEMENTS

We thank members of the Oliver Lab for technical assistance and mouse colony management (K.S. Lucas, D. Soltero, K. Spainhower, R. Uzachenko), and administrative support (X.C. Cheng, L. Houston, L. Youngblood), and Dr. Greg Wang’s lab for CUT&RUN expertise (Y. Guo). We appreciate MedGenome staff for sequencing services, and Duke Human Vaccine Institute for use of the tapestation. We acknowledge support from the NIH National Cancer Institute (NCI) under award U24CA213274 (TGO), R01-CA244841, and R01-CA187457 (TGO). T.G.O. received support as a Duke Science & Technology Scholar and from the Lung Cancer Initiative of North Carolina. K.A.O. was supported by NCI R01CA273585, P50CA70907, V Foundation (T2021-011), and the Department of Defense (DoD LC190249). L.T.I. received support from the American Cancer Society (PF-25-1366291-01-PFCBI). A.S.I. received support from the NIH NCI (F31 CA275295). We thank the BioRepository & Precision Pathology Center (BRPC), a shared resource of the Duke University School of Medicine and Duke Cancer Institute, which receives support from the P30 Cancer Center Support Grant (P30 CA014236) and from the Cooperative Human Tissue Network (UM1 CA239755) and the Duke Cancer Institute Flow Cytometry Core.

## EXPERIMENTAL MODEL AND SUBJECT DETAILS

### GEMMs

*Rb1^fl/fl^;Trp53^fl/fl^;Rbl2^fl/fl^*(RPR2)^61^, *Rb1^fl/fl^;Trp53^fl/fl^*;*LSL-Myc^T58A^* (RPM)^60^, *Sox2-Ires-GFP^LSL/LSL^;Nkx2-1^fl/fl^;Lkb1^fl/fl^*(SNL)^40^, *Lkb1^fl/fl^;Pten^fl/fl^*(LP)^18^, and *Kras^G12D/+^;Trp53^fl/fl^*(KP)^59^ mice have been previously described. Methods for tumour initiation and monitoring can be found in the listed references. Ad5-Cre with the following promoters was used: RPM (10 Ad-CGRP-Cre initiated tumours, 5 Ad-CMV-Cre initiated tumours), RPR2 (4 Ad-CGRP-Cre initiated tumours, 4 Ad-CMV-Cre initiated tumours), KP (8 Ad-CMV-Cre initiated tumours), LP (6 Ad-CMV-Cre initiated tumours), SNL (4 Ad-CMV-Cre and Ad-CCSP-Cre initiated tumours). All mice were housed following the regulations set by the Institutional Animal Care and Use Committee of the University of Utah and Duke University. At 6-10 weeks of age, anesthetized SNL mice for this study were infected by intranasal instillation with 1×10^8^ plaque forming units of Ad5-CMV-Cre or Ad5-CCSP-Cre adenovirus (University of Iowa). Viruses were administered in a Biosafety Level 2+ room according to the guidelines from the Institutional Biosafety Committee.

### Human samples

Human lung cancer tissue microarray (TMA) samples were a gift from Edward Patz, Jr. TMA samples were de-identified and only information on the histopathological subtype scoring by a trained pathologist was provided. Human TMAs from patients with lung cancer were generated following written informed consent from each patient under an Institutional Review Board-approved protocol at Duke University Medical School (Protocol ID: Pro00012914). TMA samples were reviewed by an additional pathologist to certify histopathological subtype and provide percent tumour scoring. Sections with zero percent tumour were not analyzed.

### Cell lines

The human bronchial epithelial cell line BEAS-2B was acquired from the Duke Cell Culture Facility (Sigma Catalog #95102433). The BCi-NS1.1 cells, an immortalized human airway basal cell stem/progenitor cell line, was generated at Cornell University by Dr. Ronald Crystal and provided as a gift for research purposes^87^. Both cell lines were grown in 2D adherent culture plates in Bronchial Epithelial Cell Growth Basal Medium (BEBM, Lonza Catalog # CC- 3171) with Supplements and Growth Factors (Lonza Catalog # CC-4175) along with 0.2% Penicillin-Streptomycin (BEBM media). For serial passaging, media was removed and 0.25% Trypsin-EDTA (Gibco catalog #25200056) was added to dislodge cells. HEPES Buffered Saline Solution (Catalog # CC-5024) supplemented with 15% fetal bovine serum (FBS) was added to quench the reaction. The cells were centrifuged at 250g for 5 min at 4°C. The supernatant was removed, and cells were resuspended in BEBM media before plating into tissue culture treated dishes.

Human LUSC cell lines NCI-H226 (H226) and NCI-H520 (H520) were purchased from ATCC (CRL-5826 and HTB-18 respectively). Both cell lines were grown in RPMI media (Gibco # 11875093) supplemented with 10% FBS, 1% Penicillin-Streptomycin, and 1% L-glutamine (200 mM). For serial passaging, media was removed and 0.25% Trypsin-EDTA (Gibco catalog #25200056) was added to dislodge cells. Cell culture media was used to stop the reaction and cells were plated into tissue culture treated plates.

### Organoids

After sorting from tumours (see **Harvesting tissue for organoid production or single cell sequencing**), single cell suspensions containing SNL tumour cells (NGFR^+^GFP^+^) were centrifuged at 2000 rpm for 5 min at 4°C. Supernatant was removed and cells were resuspended in Matrigel before plating. Cells were grown in 3D Matrigel domes submerged in organoid culture media (OCM) [50% DMEM/F12 media (supplemented with 10% FBS, 1% penicillin/streptomycin, 1% L-glutamine, 50ng/mL Epidermal Growth Factor (EGF) (Thermo Cat # PHG031) and 50ng/mL Fibroblast Growth Factor (FGF) (Thermo Cat # PHG0369)) + 50% L-WRN conditioned media (containing <u>W</u>nt-3A, <u>R</u>-spondin 3 and <u>N</u>oggin) + 10uM ROCK inhibitor]. L-WRN conditioned media was generated according to previously published methods^104^ using L-WRN cells purchased from ATCC (CRL-3276). For serial passaging or flow sorting, media was removed and cold DMEM media was added to wells before disruption of the Matrigel plug by pipetting. The organoid and Matrigel containing media was centrifuged at 1500 rpm for 5 min at 4°C. Supernatant was removed and organoids were resuspended in phosphate buffered saline (PBS) before centrifugation at 1500 rpm for 5 min at 4°C. Supernatant was removed and cells were resuspended in 500 uL TrypLe for 15 min at 37°C. Following vigorous pipetting, the reaction was quenched with DMEM and was centrifuged at 1500 rpm for 5 min at 4°C. Cells were then resuspended in Matrigel and plated in 24 well plates. For sorting experiments, single cell suspensions were sorted, centrifuged at 1500 rpm for 5 min at 4°C, then resuspended in Matrigel and plated in 24 well plates.

## METHOD DETAILS

### Harvesting tissue for organoid production or single cell sequencing

Mice were euthanized to harvest tumour-bearing lung tissue or tracheas. For tumour cell isolation, tumour-bearing lung tissue was removed and put into cold DMEM media (10% FBS, 1% L-glut, 1% Pen/Strep) and chopped into smaller pieces mechanically with fine scissors before further dissociation was performed using the Lung Dissociation kit (Miltenyi Biotech Cat # 130-095-927) at 37°C following the manufacturer’s instructions. Dissociated cells were filtered using a 100 µm strainer and collected into conical tubes. Cells were pelleted by centrifugation at 1500 rpm at 4°C for 5 min and then resuspended in ammonium chloride potassium (ACK) lysis buffer to lyse red blood cells. The lysis reaction was quenched with addition of PBS and cells were centrifuged. Supernatant was removed and cells from tumour-bearing lungs were resuspended in FACS buffer (PBS containing 2% FBS) for subsequent flow sorting and single cell sequencing.

### Flow sorting tumour cells

Single cell suspensions from tumour-bearing lungs or tumour organoids (see **Harvesting tissue for organoid production or single cell sequencing** and **Organoids**) were stained with a rabbit anti-NGFR (Abcam Cat #ab8875 or ATS Cat # AB- N01AP) primary antibody for 30 min on ice. Cells were then washed with FACS buffer and stained in FACS buffer containing a fluorescent secondary antibody (APC-goat anti-rabbit Invitrogen Cat # A-10931) and a fluorescent conjugated antibody (BV605-rat anti-CD45 Biolegend Cat # 103155, for tumours). Cells were then incubated on ice and in the dark for 20-30 min. Following incubation, cells were washed with FACS buffer and centrifuged. The supernatant was discarded, and cells were resuspended in FACS buffer containing DAPI to detect dead cells. Cells were then sorted using flow cytometry according to the outlined gating scheme on a cell sorter (either BD Bioscience FACSAria or Beckman Coulter MoFlo Astrios). Flow sorted tumour cells were used for single cell RNA-sequencing or for plating of organoids within 30 minutes directly following sorting (see **Single cell RNA-sequencing** or **Organoids** below). For SNL Ad-CMV-Cre tumour cells used for sequencing, GFP+ and GFP+/NGFR+ cells were sequenced in two independent experiments. For SNL Ad-CCSP-Cre tumour cells used for sequencing, GFP+ cells were used for the first sequencing (S1) and a mixture of GFP+, GFP+/NGFR+, and CD45+ cells were used for the second sequencing (S2).

Ultra-Comp eBeads Plus compensation beads (Invitrogen Cat # 01-3333-41) were used to set compensation for the NGFR primary antibody.

### Single cell RNA-sequencing

Single-cell suspensions from flow sorted GFP^+^ and NGFR^+^ GFP^+^ SNL tumours or SNL tumour organoids were prepared for sequencing according to the 10x Chromium platform protocols. (https://support.10xgenomics.com/single-cell-gene-expression). The SNL Ad-CMV- and Ad-CCSP-Cre initiated tumour samples sorted for GFP^+^, and the SNL Ad-CMV-Cre GFP^+^/NGFR^+^ tumour cells were prepared and sequenced at the Huntsman Cancer Institute High-Throughput Genomics Core at the University of Utah. The SNL Ad-CCSP-Cre mixture of GFP+, GFP+/NGFR+ (S2) tumour sample and all organoid samples were prepared in-house at Duke University and sequenced by MedGenome. Libraries were prepared with the Chromium 3’ single cell gene expression (GEX) kit V2 chemistry for Ad-CMV-GFP^+^ (10x Genomics Cat # PN-120237) and V3 chemistry for Ad-CMV-NGFR^+^GFP^+^, Ad-CCSP-GFP^+^, Ad-CCSP-NGFR^+^GFP^+^, and all organoid samples (10x Genomics Cat # PN-1000141). In brief, individual cells were barcoded with 16-bp 10x barcodes and cell-specific transcript molecules were tagged with 10bp unique molecular identifiers (UMIs), according to the manufacturer’s instructions. Reverse transcription reagents and 10x gel beads were loaded on Chromium Chip Gs (10x Genomics Cat # PN-1000136) and into the Chromium single-cell controller (10x Genomics Cat # PN-120263) to form GEMs. Within singular GEMs, cDNA generated from captured and barcoded mRNA was synthesized by reverse transcription following the manufacturer’s recommendations. Subsequent A tailing, end repair, adaptor ligation, and sample indexing were performed for each sample according to the manufacturer’s instructions. The resulting cDNA libraries were assessed on Agilent D1000 or D5000 ScreenTape on an Agilent Technology 2200 TapeStation system and quantified by quantitative PCR using a KAPA Biosystems library quantification kit (Roche Cat # 070960140001). Individual libraries were normalized and sequenced on an Illumina NovaSeq 6000, with the exception of the Ad-CMV-GFP^+^ sample that was sequenced on an Illumina HiSeq, with 10x-recommended read lengths and aiming for >20k reads per cell.

### 10x single Cell RNA-seq data processing Demultiplexing and data alignment

CellRanger was used to demultiplex and align reads per sample for downstream analysis. Single cell RNA-seq data from SNL samples were first demultiplexed with 10x cellranger mkfastq to create fastq files. For each sample, reads were then aligned to the GRCm38 (mm10) mouse genome. For SNL tumour organoids, reads were aligned to a custom mouse genome containing sequences for *fLuc*, *Venus, Cas9,* and e*GFP* (GRCm38-mm10-2020-A)^69^. Count barcodes and UMIs were generated using cellranger count.

### Quality control, data combination, and normalization

All clustering, cell type identification and quality control (QC) was performed in Python (v3.8.8) utilizing Scanpy (v1.10.0) according to current expert recommendations for single cell best practices^105^. All cells sequenced from Ad-CMV- or Ad-CCSP-Cre initiated tumours or tumour organoids were read into sparse matrix anndata objects created from filtered feature matrices with scanpy.read_ 10X_mtx(), and quality metrics were calculated using scanpy.pp.calculate_qc_metrics(). Low quality cells and potential doublets were excluded by selecting for cells with ≤15% mitochondrial content and the following criteria for respective sample types: SNL primary tumour samples with cells containing >1000 total counts and between 1000 and 6000 genes with at least one read; SNL tumour organoid samples with cells containing >1000 total counts and between 2000 and 8000 genes with at least one read. Human patient data was accessed from Wu et al^70^, patient tumours identified as squamous were used, and no additional QC filtering was performed. Head and neck patient data was accessed from Kurten et al^95^, utilizing only CD45^-^ samples and were filtered for cells containing >500 total counts and between 500 and 4000 genes with at least one read. Esophageal patient data was accessed from Zhang et al^96^, utilizing only CD45^-^ samples annotated as epithelial, and no additional QC filtering was performed. Normalized counts were calculated with scanpy.pp.normalize_total() and a target sum of 10,000. Integrated anndata objects containing multiple scRNA-seq datasets were combined using adata.concatenate() with join=’outer’.

### Batch correction and clustering

Data were further processed using Scanpy (v1.10.0) and scVI-tools (v0.17.4) and benchmarking standards to minimize batch effects while maintaining true biological variability, particularly across integrated objects^106^. Specifically, we utilized scvi.model.SCVI.setup_anndata() to initialize the integration and clustering of datasets from concatenated anndata objects containing all samples and genes, continuous covariate keys set as percent mitochondrial counts, and batch keys identifying samples prepared or sequenced at different times. The model was trained with default parameters, an early stopping patience of 20, and a maximum of 400 epochs, using the model.train() function on the top 10,000 variable genes determined using the sc.pp.highly_variable_genes() function. The latent representation of the model was obtained with model.get_latent_representation() and added to the .obsm of each anndata object. Neighbors were then calculated with sc.pp.neighbors() with use_rep set to the .obsm category added from the latent representation. UMAP embedding was performed using sc.tl.umap() with min_dist=0.5. Finally, Leiden clusters were generated with sc.tl.leiden() with resolution set to 1.0 for initial steps. As is required throughout the scVI pipeline, we utilized raw counts for all steps above. Additional rounds of quality control and data filtering were performed per dataset by assessing n_genes_by_counts, total counts, and percent mitochondrial counts per cluster. In general, the model tends to cluster low quality and doublet cells together, so clusters with exceptionally high or low average genes_by_counts, total counts, or mitochondrial content were labeled low-quality and considered for removal from the dataset. Removal was performed after assessing gene expression based on known markers of tumour and normal lung cells, and marker genes per cluster as determined by sc.tl_rank_genes_groups(), to help ensure that biological cells that normally have higher or lower n_genes_by_counts, total_counts, or percent mitochondrial content were not aberrantly filtered out. Additionally, from SNL primary tumour samples and human patient data, we identified and removed non-tumour cell types including immune, endothelial, fibroblast, and normal lung cell types using a panel of common marker genes. For the human patient data, adenocarcinoma clusters were also removed based on expression of genes known to be high in lung adenocarcinomas and not expressed in squamous tumours. Each time a cluster was removed, we ran the scVI pipeline on the new anndata object iteratively through this quality control step until there were no longer any low-quality cell clusters or non-tumour cells in the anndata object. Additionally, each time clusters were removed from a larger anndata object, the pipeline was re-run for optimal clustering. Final Leiden clusters were determined with sc.tl.leiden() resolution set to 1 for the human LUSC patient data (**Fig. 4**), 0.75 for SNL tumour organoids (**Fig. 5**), 1 for SNL Ad-CMV and Ad-CCSP cells combined (**Fig. 3**), 0.5 for SNL Ad-CMV tumour cells (**Fig. 2**), 0.5 for both head & neck and esophageal patient data (**Fig. 8**). After final leiden clustering, the embeddings determined on highly variable genes were added to anndata files that contained all genes with raw read counts. Counts were then normalized and log transformed using sc.pp.normalize_total() with a target_sum=1e4, followed by sc.pp.log1p() using default settings. UMAPs of individual genes were produced using normalized counts, and all gene scores and DEG analysis was performed on log transformed counts. Source code to reproduce analyses methods is on GitHub (https://github.com/TGOliver-Lab/Izzo_Hillock_2025) and publicly accessible upon publication.

### Calculation of basal, mucinous adenocarcinoma, tuft, and hillock cell scores and correlation analysis

The general hillock-cell score was derived by identifying genes enriched in the *Krt13^+^* cells from Montoro et al, Nature, 2018^6^ and Plasschaert et al, Nature, 2018^7^ to create a 65-gene “conserved hillock signature” (genes with conserved significant enrichment versus other normal lung cell types in both independent datasets). The “basal score”, “goblet score”, “club score”, “ciliated score” and “tuft score” are derived from droplet-based sequencing done from isolated epithelial cells from mouse trachea^6^. The top 100 differentially-expressed genes from a ranked list of differentially expressed genes (n = 1,137) from mouse mucinous adenocarcinoma tumours were used to generate the mucinous adenocarcinoma score^62^. The “basal hillock score” (n=170) was generated from differentially-expressed genes comparing basal hillock cells to normal pseudostratified lung epithelium (Log2FC>0.5) and the “luminal-hillock score” (n=335) was generated from differentially-expressed genes comparing luminal-hillock cells to normal pseudostratified lung epithelium (Log2FC>0.5)^37^ (see **Supplementary Tables 1-5**). Mouse gene scores or human gene scores were converted to their respective species orthologs using Gprofiler^107^ in Python. Hillock, basal, mucinous adenocarcinoma, basal hillock, luminal hillock, goblet, club, ciliated, and tuft scores were computed for each cell per scRNA-seq library by converting expression data to a AnnData object and utilizing the scanpy.tl.score_genes function in Scanpy on normalized and log transformed data using sc.pp.normalize_total() with a target_sum=1e4, followed by sc.pp.log1p() using default settings.

### CIBERSORTx

CIBERSORTx is a tool developed by the Alizadeh and Newman Labs at Stanford University that uses single-cell RNA-sequencing expression data input to estimate abundances of defined cell types in a bulk cell population, such as a bulk tumor^108,109^. We utilized the interactive user interface of CIBERSORTx at https://cibersortx.stanford.edu and the Cell Fraction module. A custom signature matrix file was built from a single cell reference sample file that included normalized single cell transcriptomic data from human squamous cell carcinoma tumors^70^ (n=100 randomly sampled cells per patient, n=18 patient tumors analyzed). Each cell was defined as “hillock-like” or “non-hillock-like” based on threshold expression values of the “hillock-cell score” and “basal score”, described above (hillock-like=z-score hillock_score>=2, z-score basal_score<=3). Bulk RNA-seq data from TCGA of LUSC tumors (n=538 tumor) were used as the input mixture file and subject to deconvolution with the number of permutations set to 50. Results of cell-type imputation were sorted by tumors with the highest fraction of “hillock-like” cells to the lowest fraction. Independent deconvolution of the LUSC TCGA tumors was also performed using the published leukocyte, LM22_Cibersort signature matrix^108^ to deconvolve each LUSC tumor into 22 immune cell types. Correlations of immune cell fractions (**Supplementary Table 10**) and KLF4-fraction with the KRT13-like fraction in each bulk LUSC tumor were assessed using simple linear regression tests in Graphpad Prism where CIBERSORTx graphs were also generated.

### Calculation of KLF4-up score

The “KLF4-up score” (n=51) was generated through the intersection of genes upregulated when comparing doxycycline treated to no doxycycline control of *TRE-KLF4*-expressing BEAS-2B cells (converted to mouse orthologs) and downregulated comparing *sgKlf4i-1* and *sgNTCi* SNL tumour organoids (see **Bulk RNA-seq sample and data processing from BEAS-2B and SNL tumour organoids** and **Supplementary Table 13**). The KLF4-up score was computed for each cell per scRNA-seq library by converting expression data to a AnnData object and utilizing the scanpy.tl.score_genes function in Scanpy.

### Calculation of cell cycle phase

Cell cycle analysis was performed using the scanpy.tl.scores_cell_cycle function. S-phase and G2/M-phase gene sets were derived from cc.genes in Seurat^71^ and were converted to mouse homologs. Each cell was assigned a cell cycle phase based on gene score expression using the scanpy.tl.scores_cell_cycle function. Gene scores were used for statistical comparison between assigned cell states to compare between predicted populations in S and G2/M phases.

### Cell state assignment and correlation analysis

Cell states for SNL primary tumours were determined by evaluating marker gene expression and gene signature scores in Leiden clusters. Cell state assignments were validated by gene expression per cell state and by evaluation of DEGs from each cell state compared to all other cells (see **Differential gene expression analyses**). For the human patient lung squamous cell carcinoma, esophageal squamous cell carcinoma, and head and neck squamous cell carcinoma data, cell state was assigned by determining a “hillock-like” cutoff. The z-score for the hillock score and basal score were calculated, and cells were determined to be “hillock-like” if the hillock score z-score was >=2 and the basal score z-score was <=3, which excluded highly basal-like cells while maintaining cells with high hillock gene expression compared to the overall cell population. Correlation analysis was performed using the sc.tl.dendrogram and the sc.pl.correlation_matrix functions in scanpy with default parameters using the Pearson method.

### Differential gene expression analyses

Differentially-expressed genes (DEGs) for all samples was determined using the scanpy.tl.rank_genes_groups function in Scanpy using the Wilcoxon Rank Sum Test on log transformed data. This test compares each group to all other groups to determine top DEGs in a specific cell population. ENRICHR analyses^110–112^ were performed on the top 500 upregulated genes (DEGs) in the SNL mouse tumour cell types (basal-like vs all others, basal hillock-like vs all others, and luminal hillock-like vs all others), 500 genes in the human tumour cell type (basal-like vs hillock-like), and 500 genes in organoids (basal-like vs hillock-like). For ENRICHR analysis of normal hillock genes from Montoro et al, Nature, 2018, DEGs for hillock cells were included if the log2FC>=0.5 and all *Krt13/Krt4*-enriched genes from Plasschaert et al, Nature, 2018, were included.

### Differential gene expression analyses for cell surface protein encoding genes

Differentially-expressed genes (DEGs) for all samples was determined using the scanpy.tl.rank_genes_groups function in Scanpy using the Wilcoxon Rank Sum Test. Genes were then filtered using the CSPA mouse or human cell surface protein list^113^ combined with the human in silico cell surface protein atlas (all predicted surface proteins as defined by SURFY)^114^ for human or the SURFY list converted to mouse orthologs for mouse.

### CUT&RUN sample processing and analysis

SNL tumour organoids were collected into a single cell suspension as described in **Organoids** and resuspended in OCM media. CUT&RUN was performed using the Cutana CUT&RUN protocol version 2.1 (EpiCypher). In brief, 500,000 organoid cells per CUT&RUN reaction were washed in Wash Buffer with 0.05% digitonin, incubated with activated ConA beads (EpiCypher Cat # 21-1401), and subject to antibody binding overnight (anti-KLF4, R&D Systems AF3158, 0.5 µg; anti-H3K27Ac, CST 8173S, 0.5 µg; IgG, CST 2729S, 0.5 µg). pAG-MNase was used to digest the chromatin, the released DNA fragments were purified using an NEB Monarch Spin PCR & DNA Cleanup Kit (NEB # T1135). 5 ng of DNA per reaction was used as input for library preparation using the NEBNext Ultra II DNA Library Prep Kit (NEB # E7645S). Libraries were sequenced on a NovaSeq instrument with 150 bp paired end reads targeting 5 to 10M reads per sample. We obtained over 5M reads for the anti-KLF4 CUT&RUN.

Raw sequencing read data was input into the nf-core/cutandrun pipeline (version 3.2.2) with default parameters and aligned to the mouse mm10 genome^115,116^. Peaks were called using both SEACR^117^ then were annotated using HOMER^118,119^ with the mm10 reference mouse genome. SEACR annotated peaks were filtered by protein-coding and promoter-TSS annotations before ENRICHR analysis. The standard nf-core/cutandrun .bigWig output files were used in IGV for genome track visualization.

### KLF4 modulation in organoids

For lentiviral CRISPR-mediated inhibition, the following guideRNAs targeting *Klf4* were cloned into the dCas9 KRAB vector^82^ (Addgene # *71236*) (dCas9 KRAB): sg*Klf4i–1 (5’ CGCGCCGCCGACAGCCACAG 3’), sgKlf4i–2 (5’ AGTGTCCCCCACCGTTGTCG 3’), and sgNTCi (5’ ACCTGATACGTCGTCGCGTA3’).* pLV hU6-sgRNA hUbC-dCas9-KRAB-T2a-Puro was a gift from Charles Gersbach (Addgene plasmid # 71236; RRID:Addgene_71236).

Human *KLF4* was overexpressed in tumour organoids under the control of a tetracycline response element (TRE). *KLF4* overexpression was induced in organoids using 1 µg/mL of doxycycline. Sequencing was validated and confirmed through Plasmidsaurus. For knockdown and overexpression, sequence-validated plasmids were used to generate low titer or high-titer lentivirus. 293T cells were transfected with a three-plasmid transfection system including the lentiviral vector, pCMV-dR8.2 dvpr, and pCMV-VSV-G for production of low titer virus or high titer virus.

For infection with high titer lentiviruses, organoid cells were dissociated as described for serial passaging and resuspended in 0.5mL of organoid culture media (OCM) [50% DMEM/F12 media (supplemented with 10% fetal bovine serum, 1% penicillin/streptomycin, 1% L-glutamine) + 50% L-WRN conditioned media (containing Wnt-3A, R-spondin 3 and Noggin) + 10uM ROCK inhibitor] containing polybrene (10 µg/mL) and high-titer lentivirus. Cells were then centrifuged at 300g for 30 min at room temperature within a 24 well plate. Organoid cells were then resuspended in the virus containing media and centrifuged at 2000 rpm for 5 min at 4°C. Virus containing media was removed and stored before organoid cells were resuspended in Matrigel and plated. The virus containing media was then added back to the organoid cells after Matrigel solidified. The cells were then allowed 2-3 days to grow before the media was changed to contain 1 µg/mL of puromycin or 2.5 µg/mL of blasticidin for the dCas9-Krab *Klf4* knockdown and *TRE-KLF4* overexpression constructs, respectively.

### Organoid growth assays

To determine differences in organoid growth rates, organoids were first dissociated to single cells (see **Organoids**) then passed through a 35 µm filter to ensure single cells. Organoid cells were then plated in 20 µL Matrigel domes in 48 well tissue culture plates and 200 µL of OCM media per well was added to cover the domes. Media was changed every 3-4 days and for organoids treated with doxycycline (doxy) (TRE-KLF4 and TRE-GFP organoids only), fresh doxy was added every 48 hrs. Wells were imaged daily using a 4x objective and eight images were taken per well to capture the entire Matrigel dome. Organoids were counted blindly using ImageJ and the number of organoids per well was determined by combining all organoids counted in eight images. Wells in which the entire plug was not captured in eight images were removed.

### Organoid drug treatment assays

To determine differences in organoid growth rates with drug treatments, organoids were first dissociated to single cells (see **Organoids**) then passed through a 35 µm filter to ensure single cells. Single cell suspensions were treated with cisplatin, paclitaxel, or hydrogen peroxide at the indicated doses for 15 minutes while rotating in OCM. Organoid cells were then plated in 20 µL Matrigel domes in 48 well tissue culture plates and 200 µL of OCM media per well containing drug was added to cover the domes. Media was changed after 3 days to OCM without drug. For extended cisplatin treatment experiments, on day 7 after cisplatin treatment, individual wells were treated with TrypLE to dissolve the Matrigel, organoids were spun down, then replated in 20 µL. Confluency was determined using Incucyte live cell imaging (Sartorius) using the GFP fluorescence to indicate live organoids (GFP is expressed by all cells in SNL tumour organoids). For N-acetylcysteine (NAC) rescue experiments, single cell suspensions were treated with cisplatin or vehicle for 15 minutes then plated in Matrigel. Following solidification of Matrigel, OCM media containing cisplatin or vehicle with the addition of NAC was added and incubated with cells for the first 3 days before media was changed to OCM without drug or NAC.

### Cell line growth assays and drug treatments

For NCI-H520, BEAS-2B, and BCi-NS1.1 cells, cells were plated in 96 well plates and confluency was monitored using Incucyte live cell imaging (Sartorius). For NCI-H226, cells were plated in 96 well plates and cell growth was determined compared to control conditions using the CellTiter-Glo luminescence-based ATP detection assay (Promega #G7570). For cell lines expressing *TRE-GFP* or *TRE-KLF4*, doxycycline was added on the day of plating and every 2-3 days when media was refreshed. For drug treatments, either cisplatin or paclitaxel were added 1 day after plating at the indicated doses.

### Formalin fixation of organoids

Media was removed from wells and Matrigel plugs were washed with PBS once before resuspension in 0.5 mL of PBS. Resuspend organoids were fixed using 10% formalin overnight and then transferred to 70% ethanol. Fixed organoids were then centrifuged at 2500 rpm and washed with PBS twice before being placed in 2.5% agarose for embedding. Agarose embedded organoids were then placed in 70% ethanol and submitted for embedding in paraffin.

### KLF4 modulation in cell lines

For lentiviral CRISPR-mediated knockout, the following guideRNAs targeting *KLF4* were cloned into the lentiCRISPRv2 ((Addgene # 52961) (LCV2): sg*KLF4-*2 (5’-TCCCGGCCGCCCCTATGACC-3’) and sg*KLF4-3* (5’- AATTGGAGAGAATAAAGTCC-3’). For overexpression of KLF4, lentiviral infection of *TRE-KLF4* was performed.

### T7 surveyor assay

Genomic DNA was extracted following the protocol in the Qiagen DNeasy Blood and Tissue Kit (Cat # 69504). Following extraction, the targeted genomic regions were PCR amplified with the following sequence primers:

To isolate the region surrounding the guide binding site for LCV2 *sgKLF4-2*, the following primers were used LCV2 *sgKLF4-2* FWD Primer-1: 5’ ACCAAGAGCCACTGAACGAG 3’; LCV2 *sgKLF4-2* REV Primer-1: 5’ CCGGATCGGATAGGTGAAGC 3’; LCV2 *sgKLF4-2* FWD Primer-2: 5’ GTGTATGCCCGTGGTGCGA 3’; LCV2 *sgKLF4-2* REV Primer-2: 5’ GATGGGTCAGCGAATTGGAGA 3’. The expected band sizes were 620 bp and 300 bp for *sgKLF4-2* primer set 1, and 550 bp and 170 bp for *sgKLF4-2* primer set 2.

The PCR conditions used for both primer sets were 30 sec for 98°C, 35 cycles of 5 seconds for 98°C, 10 seconds for 69°C, and 20 seconds for 72°C, 2 minutes for 72°C then held at 4°C.

To isolate the region surrounding the guide binding site for LCV2 *sgKLF4-3*, the following primers were used LCV2 *sgKLF4-3* FWD Primer-1: 5’ ATCTTTCTCCACGTTCGCGT 3’; LCV2 *sgKLF4-3* REV Primer-1: 5’ TTGATGTCCGCCAGGTTGAA 3’; LCV2 *sgKLF4-3* FWD Primer-2: 5’ CCAGCACGTCAGTATGTCGG 3’; LCV2 *sgKLF4-3* REV Primer-2: 5’ CGCGTAATCACAAGTGTGGG 3’. The expected band sizes were 1000 bp and 200 bp for *sgKLF4-3* primer set 1, and 250 bp and 1000 bp for *sgKLF4-3* primer set 2.

Q5 high fidelity 2x master mix (NEB Catalog # M0492) was used for the PCR reactions according to the manufacturer protocol. PCR products were then mixed with 6x loading dye and ran on 2% agarose gels. Gels were imaged to confirm correct PCR product size. The PCR products were then gel extracted using the QIAquick gel extraction kit (Qiagen #28706). DNA concentrations were measured on a Biotek Synergy HT plate reader.

200ng of gel-purified DNA was used for endonuclease reaction. The T7 surveyor assay was performed according to the “Determining Genome Targeting Efficiency using T7 Endonuclease I” protocol from NEB. Briefly, 200ng of gel-purified DNA was hybridized in a thermocycler by initial denaturing at 95°C for 5 minutes followed by annealing from 95°C to 25°C. Annealed DNA was then subjected to the T7 endonuclease at 37°C for 15 min. Following the enzymatic reaction DNA was run on 2% gels and imaged.

### Immunoblot

Total protein lysates were prepared as previously described, separated via SDS-PAGE, and transferred to a PVDF membrane^120^. Membranes were blocked for 1 h in 5% milk followed by overnight incubation with primary antibodies at 4°C. All primary antibodies were diluted 1:1000 and include: KRT13 (Abcam Cat # 92551); KRT5 (Biolegend Cat #905501); KLF4 (CST Cat # 4038s); CAS9 (Active Motif Cat # 61798); HSP90 (CST Cat # 4877); P21 (CST Cat # 2947s); PITX1 (Novus NBP1-88644); and GFP (CST Cat #2956s). Membranes were washed for three times for 5 min at room temperature in TBS-T. Mouse and rabbit HRP-conjugated secondary antibodies (1:5000; Jackson ImmunoResearch Cat # 711-035-152 and Cat # 115-035-205) were incubated for 1 hr in 5% milk at room temperature followed by washing three times for 5 minutes at room temperature in TBS-T. Membranes were exposed to WesternBright HRP quantum substrate (Advansta) and detected on Hyblot CL film (Denville Scientific, Inc.).

### Immunohistochemistry (IHC)

Tissue from autochthonous mouse models was prepared by harvesting lungs inflated with 1X PBS. Individual lobes were separated and fixed in 10% neutral buffered formalin for 24 h at room temperature (RT), washed in PBS and transferred to 70% EtOH. Formalin-fixed paraffin embedded (FFPE) tissue was sectioned at 4–5 μm. For staining, tissue were dewaxed, rehydrated, and subjected to high-temperature antigen retrieval by boiling for 20 min in 0.01 M citrate buffer at pH 6.0 in a pressure cooker. Slides were quenched of endogenous peroxide in 3% H_2_O_2_ for 15 min and blocked in 5% goat serum in PBS/0.1% Tween-20 (PBS-T) for 1 h. Slides were incubated with primary antibodies in blocking buffer (5% goat serum or SignalStain antibody diluent, Cell Signaling Technology, Cat #8112). Slides were washed 3x with PBS-T then incubated for 1 hr with secondary antibody. For all non-CST antibodies, the secondary antibody was HRP-conjugated (Vector Labs) used at a 1:200 dilution in PBS-T. Alternatively, CST primary antibodies were detected using 150 μL of SignalStain Boost IHC Detection Reagent (CST cat#8114). DAB staining was then performed, and slides were coverslipped with the automated Epredia ClearVue coverslipper. The primary antibodies include: 1:250 KRT13 (Abcam Cat # 92551); 1:2000 KRT5 (BioLegend Cat # 905501); 1:500 HNF4α (CST Cat # 3113). For manual H-score quantification, whole slides were scanned using a Pannoramic Midi II automatic digital slide scanner (3DHistech). IHC quantification included all SNL tumours from all collected lung lobes, including squamous and adenocarcinoma tumours. H-score was determined by rating staining intensity per antibody on a scale from 0-3 then multiplying each intensity by the percentage of tumour cells that express the marker. For example, a tumour with 50% positive cells with high intensity of 3 and 20% positive cells with moderate intensity of 2 has a 190 H-score.

For human tissue TMA IHC, staining was performed as described above. H-scoring was performed on the entire TMA spot. Any spots scored as containing no tumour were excluded from analysis.

### Immunofluorescence

FFPE lung tissue (see **IHC**) and organoids (see **Formalin fixation of organoids**) were prepared as described earlier. For staining, tissue was dewaxed, rehydrated, and subjected to high-temperature antigen retrieval by boiling for 20 min in 0.01 M citrate buffer at pH 6.0 in a pressure cooker. Slides were blocked in 5% goat serum in PBS/0.1% Tween-20 (PBS-T) for 1 hr or if mouse primary antibodies were used, slides were blocked in blocking agent from the M.O.M. kit (Vector Labs Cat # BMK-2202) for 1 hr. Slides were washed 3x with PBS-T then incubated for 1 hr with secondary antibody in the dark. Then slides were washed with PBS-T and incubated for 1 hr with DAPI in PBS-T before slides were coverslipped using Fluoromout-G (SouthernBiotech Cat # 0100-01). The primary antibodies include: 1:250 KRT13 (Abcam Cat # 92551); 1:400 TRP63 (R&D Systems Cat # AF1916); 1:100 P40 (Biocare Cat # ACI3066); 1:200 KI67 (BD Cat # 556003); 1:400 NGFR (CST Cat # 8238); 1:500 KLF4 (R&D System Cat # AF3158); 1:200 LGALS3 (CST Cat # 89572). Quantification of percent positive cells was performed on non-overlapping images using CellProfiler or through blinded manual counting^121^.

### Bulk RNA-seq sample and data processing from GEMMs

5 µm serial sections of FFPE tissue from LP and KP tumour bearing lungs, initiated with Ad-CMV-Cre, were stained with H&E and Aniline blue. H&E-slides were used to confirm histology and guide bordering of Aniline-blue-stained serial sections. Using a microdissection scope, bordered tissue samples were dissected from consecutive serial sections using sterile scalpels and razors. 10-14 pieces of each sample isolated were pooled in a microcentrifuge tube containing 100% ethanol. RecoverAll Total Nucleic Acid Isolation Kit for FFPE (Thermo Fisher Cat # AM1975) was used to isolate RNA from FFPE tissues according to the manufacturer’s protocol. RNA isolation from ∼15 mg flash-frozen SNL (CMV-Cre initiated), RPM (10 CGRP-Cre initiated tumours, 5 Ad-CMV-Cre imitated tumours), and RPR2 (4 CGRP-Cre initiated tumours, 4 Ad-CMV-Cre initiated tumours) primary tumours was performed using the RNeasy Mini Kit (Qiagen Cat # 74104) with the standard protocol. RNA from SNL, LP, KP, RPM, and RPR2 tumours were subject to library construction with the Illumina TruSeq Stranded mRNA Sample Preparation Kit (Cat # RS-122-2101, RS-122-2102) or the Illumina TruSeq Stranded Total RNA Library Ribo-Zero Gold Prep kit (Cat # RS-122-2301), according to the manufacturer’s protocol. LP and KP tumour samples were sequenced on an Illumina HiSeq 2500 instrument, 1×50 bp read lengths. RPR2, RPM, and SNL samples were sequenced on the Illumina NovaSeq 6000, 2×50 bp read lengths or the Illumina HiSeq 2500, 1×50 bp read lengths.

Fastq raw count files were aligned in the R statistical environment. The mouse GRCm38 FASTA and GTF files were downloaded from Ensembl release 94 and the reference database was created using STAR^122^ with splice junctions optimized for 50 base pair reads and in two pass mode to output a BAM file sorted by coordinates. Mapped reads were assigned to annotated genes in the GTF file using featureCounts version 1.6.3^123^. The output files from cutadapt, FastQC, Picard CollectRnaSeqMetrics, STAR and featureCounts were summarized using MultiQC^124^ to check for any sample outliers. To remove sources of unwanted variation from tumour RNA-seq sample preparation, all non-coding features, histones, and ribosomal RNAs were removed from count matrices for downstream analyses. The featureCount output files were combined into a single raw count matrix. Log2(counts+1)-transformed, normalized intensity values were obtained following DESeq2 analysis in R to obtain gene expression levels for GEMM tumours.

### Bulk RNA-seq sample and data processing from BEAS-2B

Total RNA was extracted from BEAS-2B cells with TRE-KLF4 after 48 hours of 250 ng/mL doxycycline treatment using a RNeasy mini kit (Qiagen Cat. No. 74104). RNA was processed by Medgenome and cDNA libraries were prepared using the Illumina TruSeq Stranded Total RNA kit. Libraries were sequenced on a NovaSeq 600/ X Plus with a read length of 100 aiming for at least 30M paired reads per sample. Each sample had over 70M reads and reads were aligned to the reference human genome (GRCh38/hg38) using STAR, read counts per gene were estimated using HTSeq, and read counts were normalized using DESeq2. Log2FC of differentially expressed genes between doxycycline and no doxycycline were used to perform GSEA using the python package GSEApy^125^ and to perform ENRICHR analysis.

### Bulk RNA-seq sample and data processing from SNL tumour organoids

Total RNA was extracted from SNL tumour organoids after digestion to single cell suspension using a RNeasy mini kit (Qiagen Cat. No. 74104). RNA was processed by the Duke Genomics Core and cDNA libraries were prepared using the Watchmaker mRNA library prep kit (Watchmaker Genomics 7BK0001). Libraries were sequenced on a NextSeq 1000 with a read length of 100 aiming for at least 30M paired reads per sample. Reads were aligned to a custom mouse genome containing GFP (GRCm38) using STAR, read counts per gene were estimated using the summarizeOverlaps function in the GenomicAlignments package of Biocondutor. Log2FC of differentially expressed genes between *sgNTCi* and *sgKlf4i-1* were used to perform GSEA using the python package GSEApy^125^ and to perform ENRICHR analysis.

### MicroCT imaging

SNL mice were imaged to monitor tumour development using microCT imaging beginning four weeks after tumour initiation by Cre-mediated recombination. Mice were anesthetized with isoflurane and imaged using a small animal Quantum GX2 microCT (Perkin Elmer). Quantum GX2 images were acquired with 2-minute scans at 45 mm resolution, 90 kV, with 88 mA of current.

## Material availability

Human LUSC and LUAD tissue used in this study are not available due to scarcity. Tumour derived organoids have limitations on availability due to derivation from primary tumours but may be made available upon reasonable request.

## Data availability

All raw and processed data have been made available through the Gene Expression Omnibus (GEO) under the superseries GSE291730. Human lung squamous cell carcinoma data from Wu et al^70^ can be found at GEO under the accession code GSE148071, HNSCC data from Kurten et al^95^ can be found at GEO under the accession code GSE164690, and ESCC data from Zhang et al^96^ can be found at GEO under the accession code GSE160269.

## Code availability

All original code has been deposited on GitHub (https://www.github.com/TGOliver-lab/Izzo_Hillock_2026) and is present on Zenodo (https://doi.org/10.5281/zenodo.18260411) which will be available at the time of publication. Any additional information required to reanalyze the data reported in this paper is available from the lead contact upon request.

## Statistical analyses

Remaining statistical analysis were performed using GraphPad Prism or Python. Error bars show mean ± SD unless otherwise specified. Significance was determined by Student’s two-tailed unpaired t-tests with 95% confidence intervals and p values <0.05, by Mann-Whitney U-test, by one-way ANOVA correcting for multiple comparisons using Tukey’s Honest Significant Difference test, by two-way ANOVA with a Sidak correction for multiple comparisons, or by Kruskal-Wallis test followed by the Dunn’s test with Bonferroni correction for multiple comparisons. All statistical details are further described in respective figure legends.

## SUPPLEMENTARY TABLES

Supplementary Table 1: Hillock score genes for mouse and human, related to Fig. 3.

Supplementary Table 2: Basal, tuft, goblet, ciliated, and club score genes, related to Fig. 3.

Supplementary Table 3: Mucinous adenocarcinoma score genes for mouse and human, related to Fig. 3.

Supplementary Table 4: Basal hillock score genes for mouse and human, related to Fig. 3.

Supplementary Table 5: Luminal hillock score genes for mouse and human, related to Fig. 3.

Supplementary Table 6: Differentially expressed genes in SNL tumour cells derived from scRNA-seq data, related to Fig. 4 and Extended data Fig. 4.

Supplementary Table 7: Reactome pathway enrichment in luminal hillock-like SNL primary tumour cells, related to Fig. 4.

Supplementary Table 8: Human patient data hillock-like cells per patient, related to Fig. 4.

Supplementary Table 9: Differentially expressed genes in hillock-like human tumour cells derived from scRNA-seq data, related to Fig. 4.

Supplementary Table 10: Immune cell population correlation to the hillock-like cell state by CIBERSORTx deconvolution, related to Extended Data Fig. 4.

Supplementary Table 11: Differentially expressed genes in hillock-like tumour organoid cells derived from scRNA-seq data, related to Extended Data Fig. 5.

Supplementary Table 12: ENCODE and ChEA Consensus TFs from ChIP-X from mouse and human hillock-like and basal-like cells, related to Extended Data Fig. 6.

Supplementary Table 13: Differentially expressed genes in BEAS-2B cells with TRE-KLF4 and SNL tumour organoids with sgKlf4i-1, related to Fig. 7.

Supplementary Table 14: SEACR peaks annotated using HOMER from KLF4 CUT&RUN in SNL tumour organoids, related to Fig. 7.

Supplementary Table 15: Differentially regulated genes encoding for cell surface proteins, related to Fig. 8.

---

**Extended Data Fig. 1.**
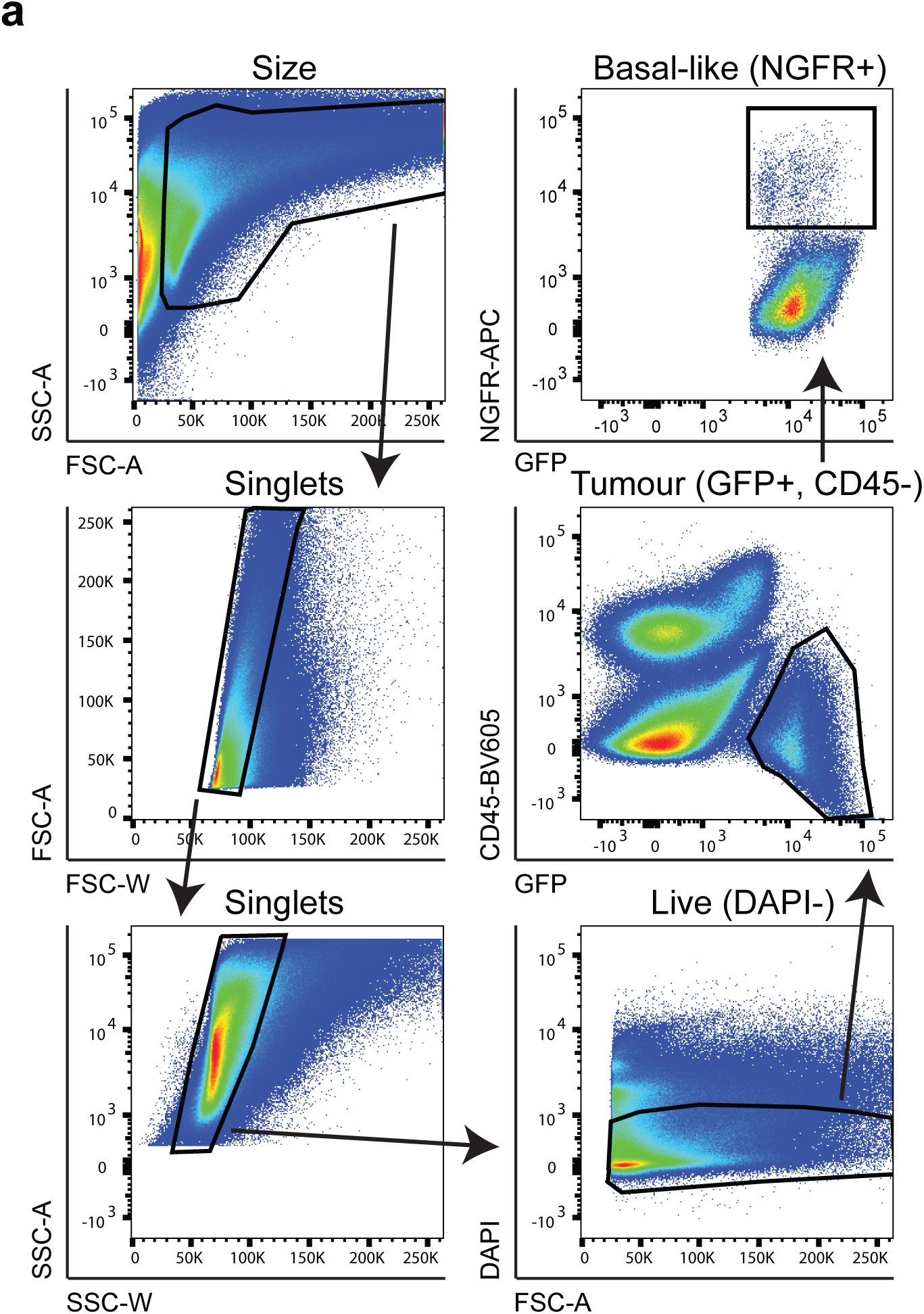
NGFR^+^ tumour cells sorted from SNL tumours for scRNA-seq, related to Fig. 2. **a)** Representative flow cytometry plots for gating strategy of DAPI^-^CD45^-^GFP^+^NGFR^+^ SNL tumour cells. SSC: side scatter, FSC: forward scatter, A: Area, W: Width. Arrows in upper left graph flow down to the right, and up back to the upper right panel, indicating progressive gating during the flow sort.

**Extended Data Fig. 2.**
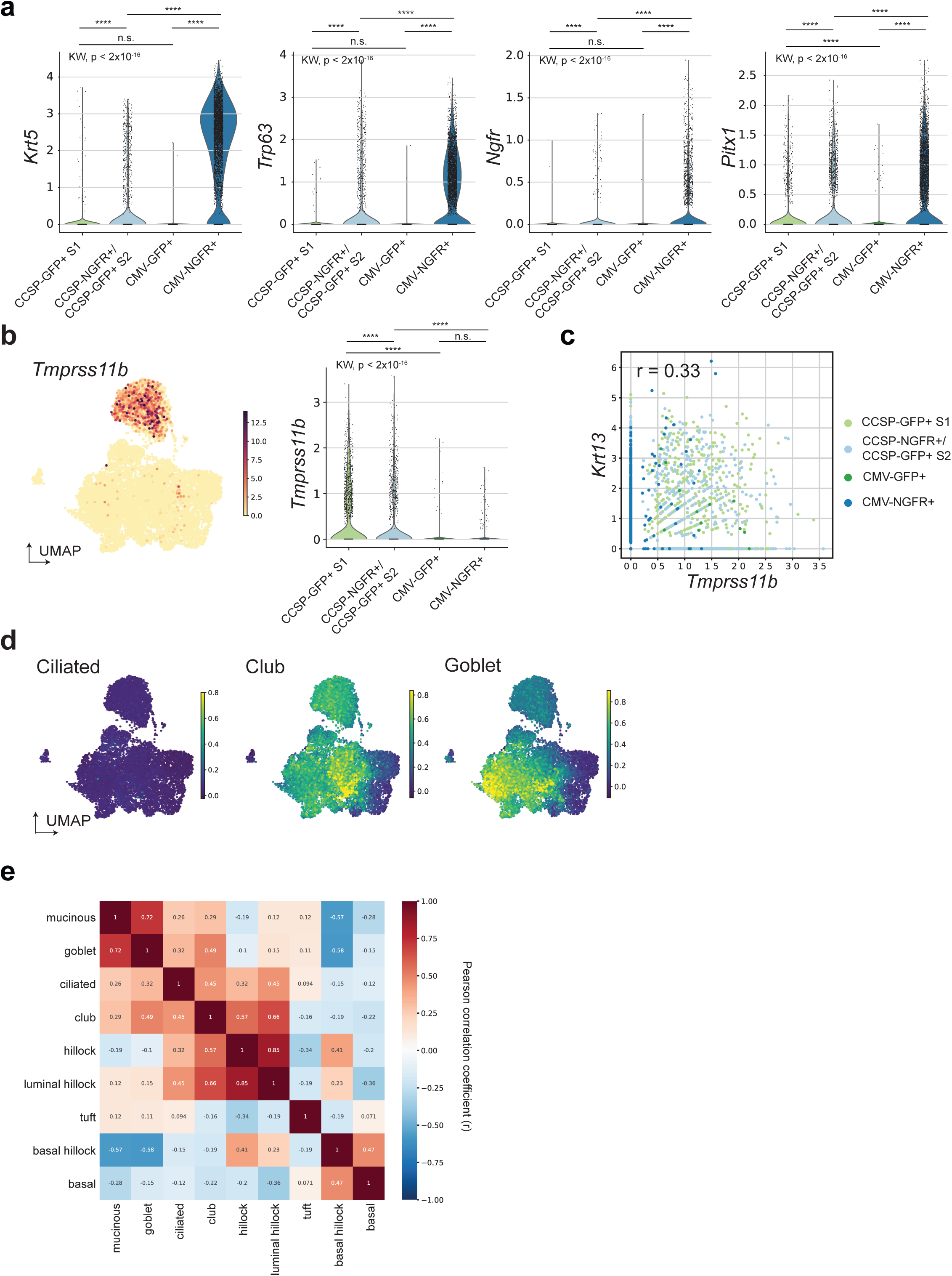
*Tmprss11b* correlates with *Krt13 expression*, related to Fig. 3. **a)** Violin plots for basal marker genes by sample. Kruskal-Wallis followed by Dunn’s test with Bonferroni correction, not all comparisons shown. **b)** scRNA-seq expression of *Tmprss11b* projected on UMAP space from cells in **3a** (left) and violin plot for *Tmprss11b* expression by sample (right). Kruskal-Wallis followed by Dunn’s test with Bonferroni correction, not all comparisons shown. **c)** Correlation between *Krt13* and *Tmprss11b* expression in cells from **3a** with the Pearson correlation coefficient (r) shown. **d)** Gene scores applied to SNL tumour cells projected onto UMAP space from **3a** (see **Methods** and **Supplementary Table 2**). **e)** Correlation analysis between gene scores in SNL tumour single cell data in **3a** with the Pearson correlation coefficient (r) shown. **For all panels,** ****p ≤ 0.0001.

**Extended Data Fig. 3.**
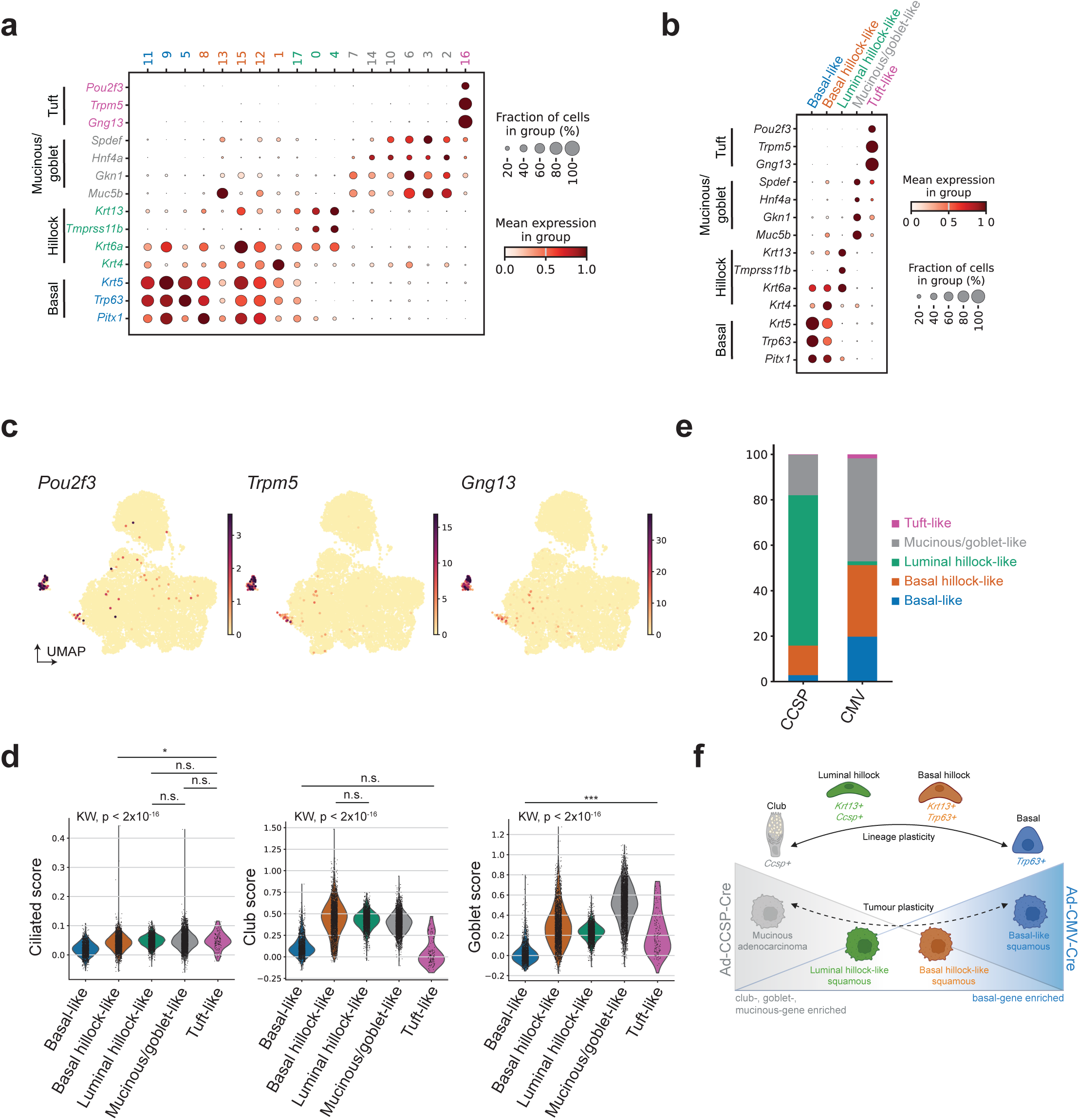
Defining cell states by transcriptional profiles in SNL tumours, related to Fig. 3. **a)** Scanpy dot plot of scRNA-seq gene expression by Leiden clustering. Text coloring of Leiden clusters indicates cell type assignment, reflected in b. **b)** Scanpy dot plot of scRNA-seq gene expression by assigned cell type. For **a)** and **b)**, Dot size indicates percent of cells expressing each transcript and the color indicates average expression per group, as indicated in figure. **c)** scRNA-seq expression of tuft marker genes projected on UMAP from cells in **3a**. **d)** Violin plots of gene score expression by cell type assignment. Kruskal-Wallis followed by Dunn’s test with Bonferroni correction, all comparison p < 0.0001 unless otherwise shown. **e)** Stacked bar plot showing percent of cells occupying each cell state in Ad-CCSP- versus Ad-CMV-initiated tumour samples from scRNA-seq data in **3h**. **f)** Schematic depicting potential lineage plasticity in normal cell types and SNL tumor cell states. Created with Biorender.

**Extended Data Fig. 4.**
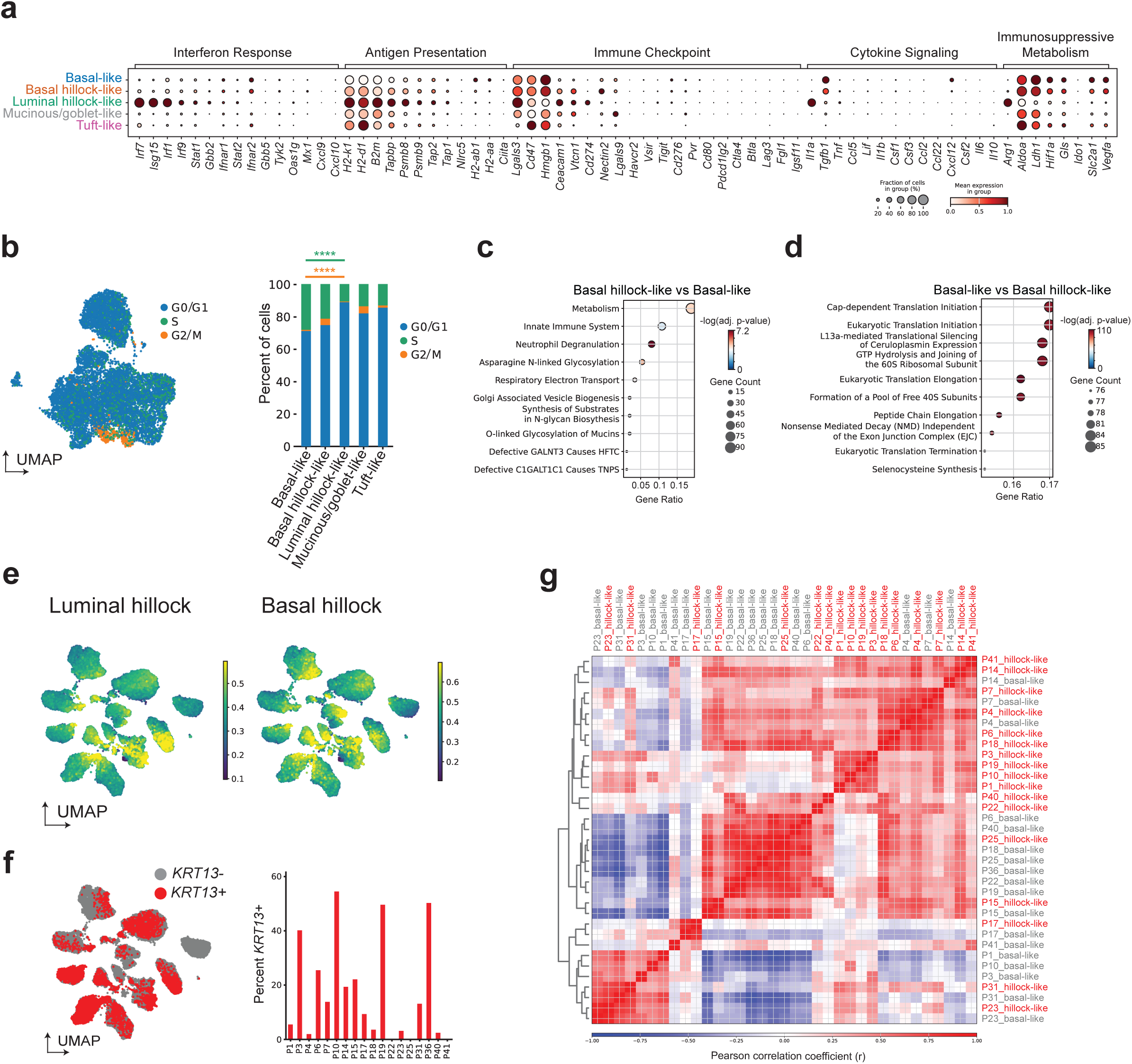
The hillock state in squamous tumours may be immunomodulatory and is relatively non-proliferative, related to Fig. 4. **a)** Scanpy dot plot of scRNA-seq gene expression of SNL tumours depicted in Fig. 3a, by Leiden clustering. Dot size indicates percent of cells expressing each transcript and the color indicates the average expression per cluster indicated in figure. Text coloring indicates cell type assignment. **b)** Cell cycle assignments applied to cells in UMAP space as in **3a** (see **Methods**) and stacked bar plot showing percent of cells occupying each cell cycle phase by cell state assignment as in **3h**. Statistics shown compare G2/M (orange) or S-phase (green) gene scores between assigned cell states. Kruskal-Wallis test followed by Dunn’s test, not all comparisons shown. **c-d)** Gene set enrichment analysis comparing basal-like and basal hillock-like SNL tumour cells from **3h**. The ENRICHR gene set used was ‘Reactome Pathways 2024’. **e)** Luminal hillock and basal hillock scores applied to human lung squamous cell carcinoma tumours in UMAP space as in **4d** (see **Methods** and **Supplementary Tables 4 and 5**). **f)** UMAP showing assignment of *KRT13* state from **4d** (left) and *KRT13* assignment per patient tumour (n=18) as a percentage of all tumour cells per sample (right). **g)** Correlation analysis of the whole trasncriptome of patient tumours divided by cell state from single cell data in **4d, j** with the Pearson correlation coefficient (r) shown. Red labels indicate hillock-like cells from each patient; grey labels indicate basal-like cells from each patient. **See also Supplementary Table 8 and 9.**

**Extended Data Fig. 5.**
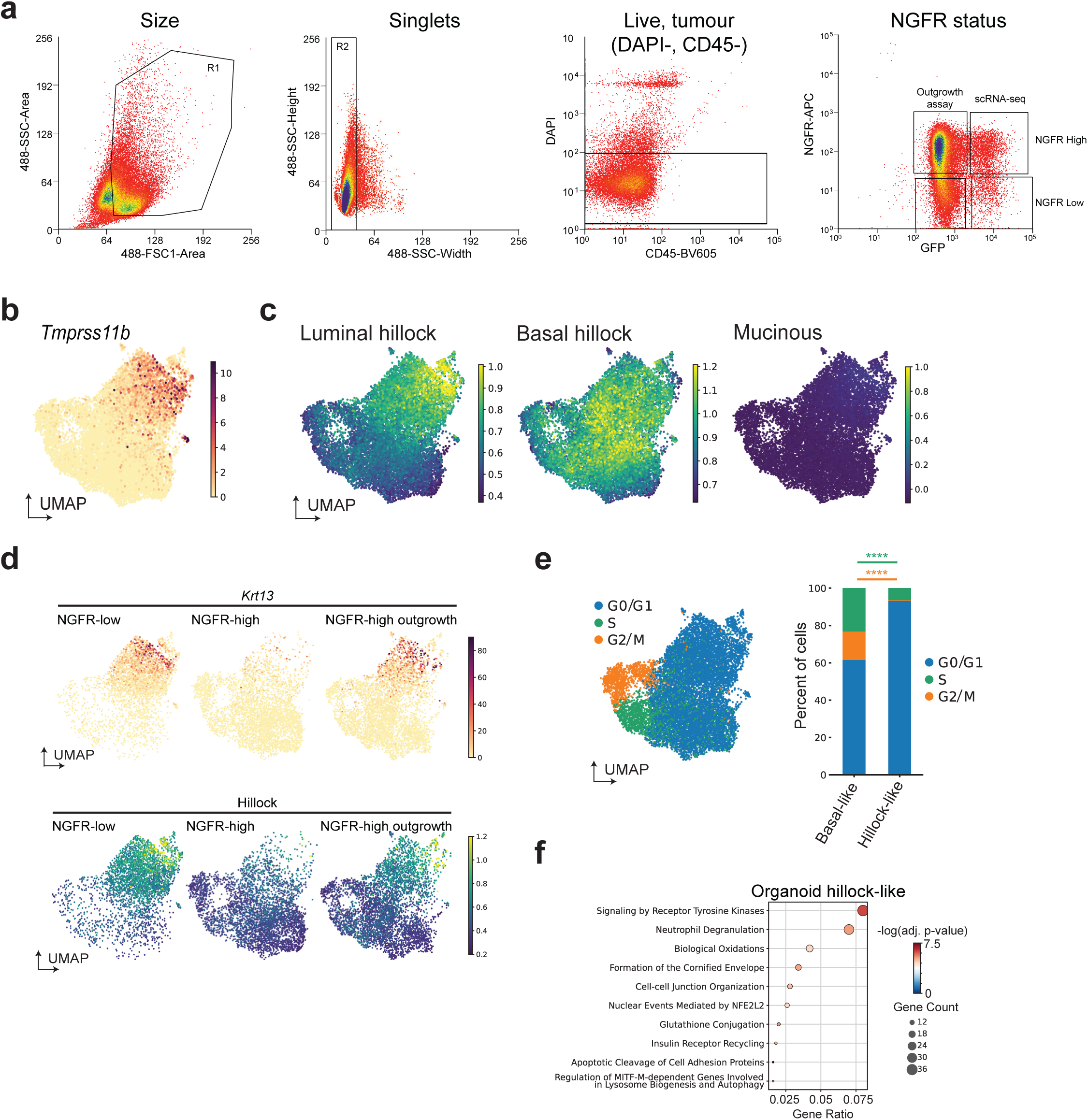
NGFR^+^ cells give rise to KRT13^+^ hillock-like cells, related to Fig. 5. **a)** Gating strategy for NGFR-high and low cells from dissociated SNL tumour organoids cells in **5f-l**. GFP/NGFR-high cells were used for the “NGFR-high” scRNA-seq sample and replated for the “NGFR-high outgrowth” scRNA-seq. A 50:50 mixture of GFP-high:GFP-low NGFR-low cells were used for the “NGFR-low” scRNA-seq sample. GFP low samples were used for organoid outgrowth assays. **b)** *Tmprss11b* expression projected onto UMAP space from **5h**. **c)** Luminal hillock, basal hillock, and mucinous scores applied to cells projected onto UMAP space from **5h** (see **Methods** and **Supplementary Tables 3-5**).**d)** Split UMAPs by sample showing *Krt13* and hillock score expression in cells from **5h**. **e)** Cell cycle assignments applied to cells projected onto UMAP space from **5h** (see **Methods**) and stacked bar plot showing percent of cells occupying each cell cycle phase by cell state assignment as in **5h**. Statistics shown compare G2/M (orange) or S-phase (green) gene scores between assigned cell states. Mann-Whiteney U-Test. ****p ≤ 0.0001. **f)** Gene set enrichment analysis comparing organoid cells in **5l** by assigned cell type. The ENRICHR gene set used was ‘Reactome Pathways 2024’. **See also Supplementary Table 11.**

**Extended Data Fig. 6.**
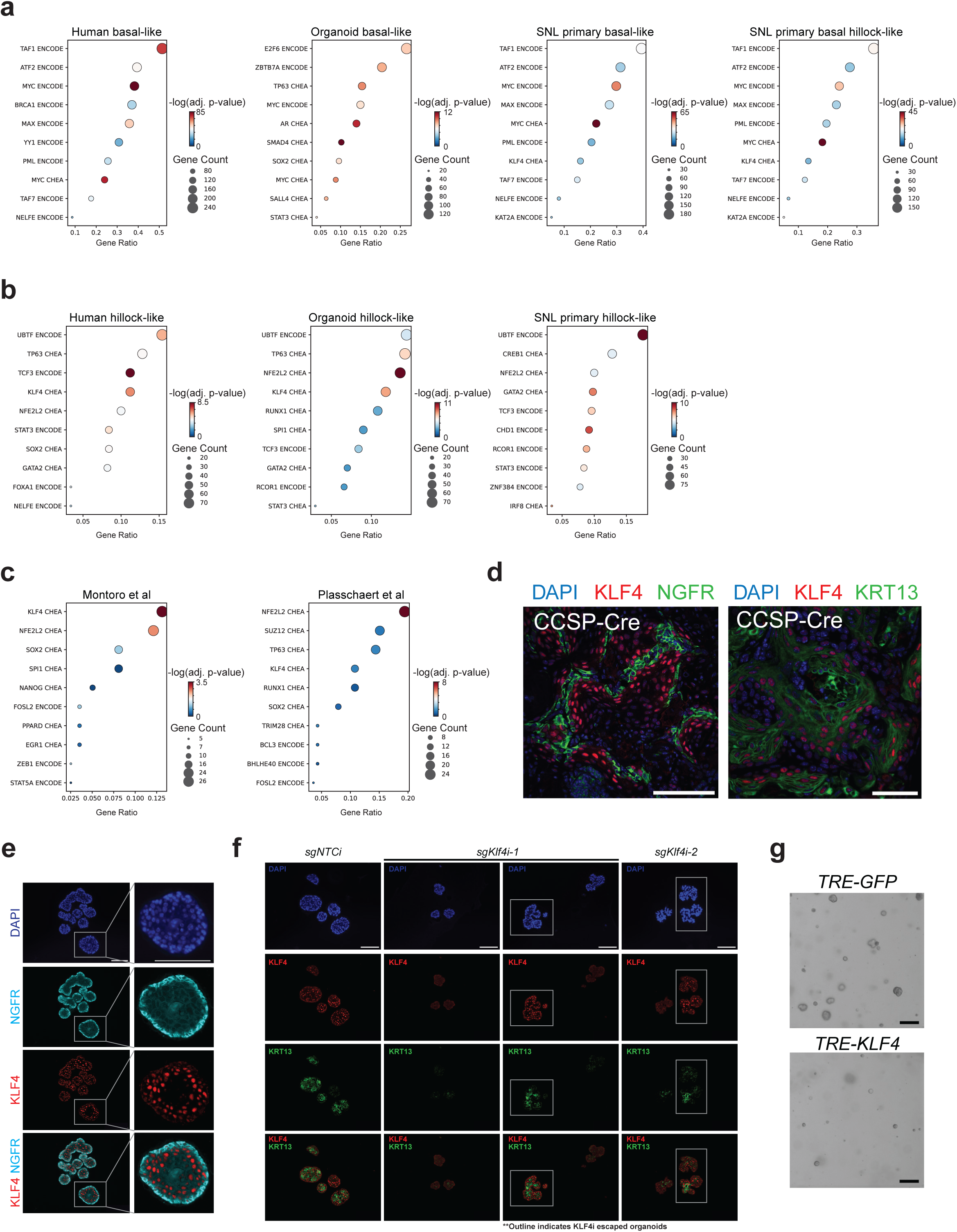
Identifying candidate regulators of the hillock cell state and KLF4 as a regulator of KRT13, related to Fig. 6. **a)** ENRICHR analysis for predicted regulators of the basal-like state in human basal-like cells, organoid basal-like cells, SNL tumour basal-like and basal hillock-like cells using ‘ENCODE and ChEA Consensus TFs from ChIP-X’. **b)** ENRICHR analysis for predicted regulators of the hillock-like state in human hillock-like cells, organoid hillock-like cells, and SNL tumour luminal hillock-like cells using ‘ENCODE and ChEA Consensus TFs from ChIP-X’. **c)** ENRICHR analysis for predicted regulators of the hillock state in normal lung tissue from the indicated publications (Montoro et al, Nature, 2018; Plasschaert et al, Nature, 2018) using ‘ENCODE and ChEA Consensus TFs from ChIP-X’ (see **Methods - Differential gene expression analyses** and **Supplementary Table 11**). **d)** IF staining for NGFR (green) and KLF4 (red) or KRT13 (green) and KLF4 (red) in squamous tumours from Ad-CCSP-initiated SNL tumours. Representative of n = 3 mice. Scale bars, 100 µm. **e)** Representative IF co-staining for KLF4 (red) and NGFR (cyan) in SNL tumour organoids. Scale bars, 100 µm. Representative of n = 2 experiments at with least 4 different 20x fields per experiment. **f)** Representative IF co-staining for KRT13 (green) and KLF4 (red) in SNL tumour organoids with KLF4 knockdown using CRISPRi. Scale bars, 100 µm. Images are uncropped coinciding to representative images shown in **6h**. White boxes indicate *sgKlf4i* expressing organoids with escaped KLF4 expression. DAPI (blue) marks nuclei. **g)** Representative brightfield images of SNL tumour organoids infected with TRE-GFP or -KLF4 treated with 1µg/mL doxycycline every 48 hrs for 7 days. Scale bars, 200 µm.

**Extended Data Fig. 7.**
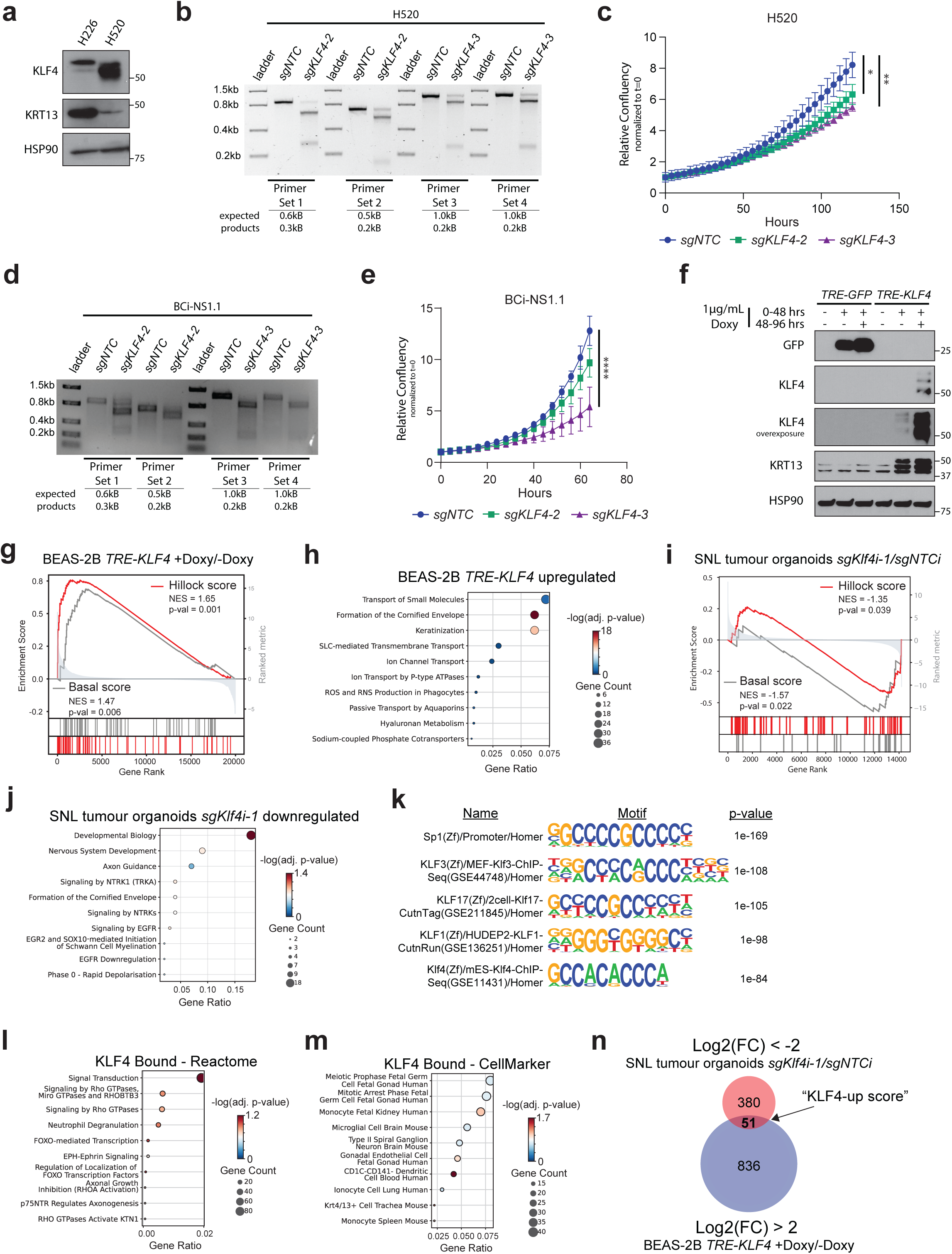
KLF4 regulates the KRT13^+^ hillock-like-state, related to Fig. 7. **a)** Immunoblot for indicated proteins in the human LUSC cell lines H226 and H520. **b)** T7 surveyor assay for *KLF4* gene editing in human H520. Expected fragment sizes after T7 cutting of mismatched (edited) DNA are shown below. **c)** Cell growth determined by confluency normalized to initial plating density of H520 cells. One-way ANOVA at endpoint. **d)** T7 surveyor assay for *KLF4* gene editing in human BCi-NS1.1 cells. Expected fragment sizes after T7 cutting of mismatched (edited) DNA are shown below. **e)** Cell growth determined by confluency normalized to initial plating density of BCi-NS1.1 cells. One-way ANOVA. **f)** Immunoblot for indicated proteins in BEAS-2B cells infected with TRE-GFP or KLF4 treated with (+) or without (-) doxycycline (doxy) for the indicated times. HSP90 serves as loading control. **g)** GSEA for the general hillock score and basal score derived from RNA-sequencing of TRE-KLF4 BEAS-2B cells treated +/- doxy as in **7g**. **h)** Gene set enrichment analysis of the top 100 upregulated genes by TRE-KLF4 in BEAS-2B cells treated +/- doxy as in **7g**. The ENRICHR gene set used was ‘Reactome Pathways 2024’. **i)** GSEA for the general hillock score and basal score derived from RNA-sequencing of SNL tumour organoids as in **7h**. **j)** Gene set enrichment analysis of the top 100 downregulated genes by *sgKlf4i-1* in tumour organoids as in **7h**. The ENRICHR gene set used was ‘Reactome Pathways 2024’. **k)** HOMER motif analysis of SEACR called peaks from KLF4 CUT&RUN in SNL tumour orgnaoids. **l,m)** ENRICHR analysis of the top 500 peaks annotated as protein coding promoters bound by KLF4 in SNL tumour organoids. The ENRICHR gene set used was ‘Reactome Pathways 2024’ and ‘CellMarker 2024’ respectively. **n)** Overlap of top genes downregulated by KLF4 knockdown in organoids and KLF4 overexpression in BEAS-2B. 51 overlapping genes used to generate the “KLF4-up” score. **See also Supplementary Table 13 and 14.**

**Extended Data Fig. 8.**
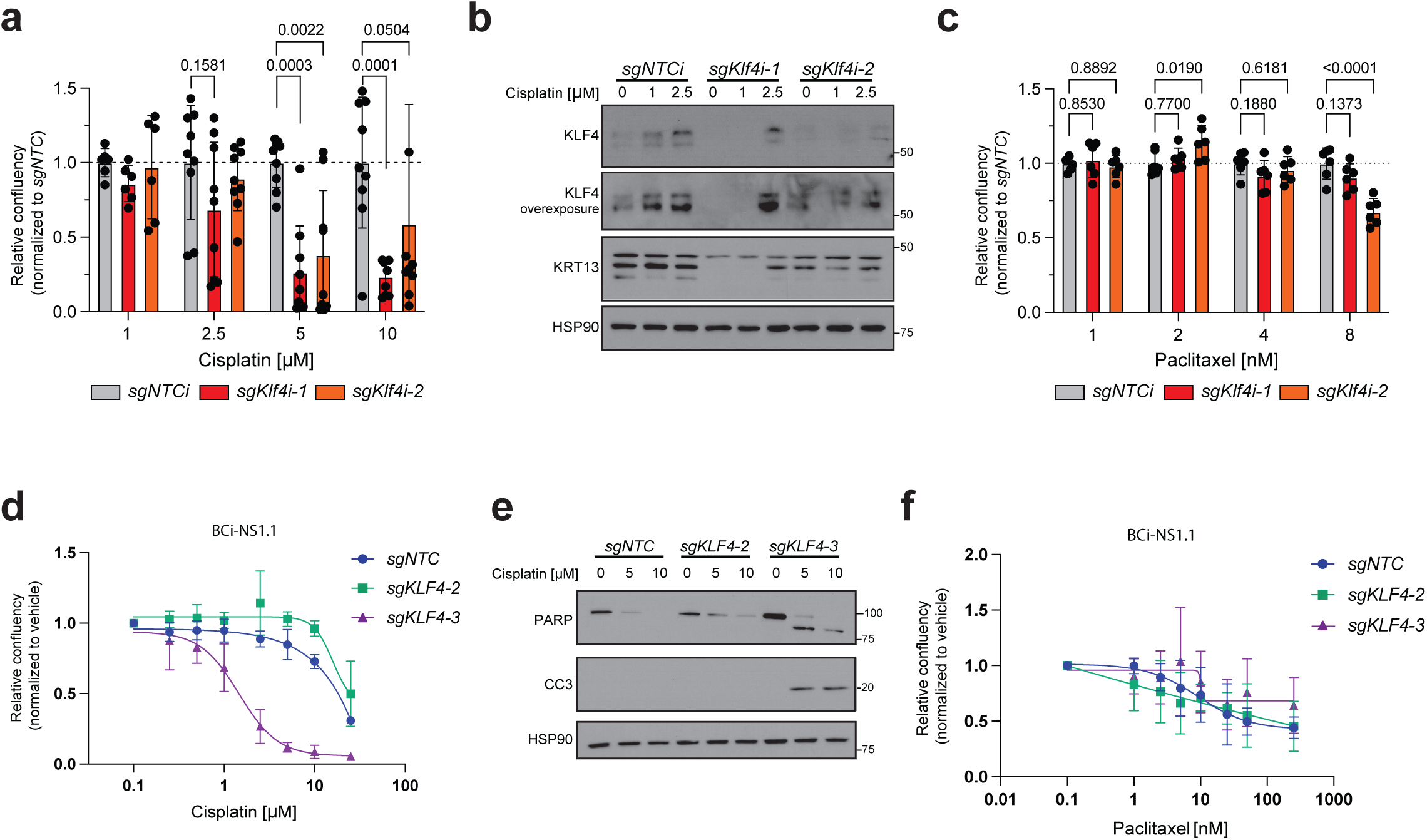
KLF4 regulates resistance to cisplatin but not paclitaxel, related to Fig. 7. **a)** Organoid growth after treatment with cisplatin determined by GFP+ confluency using an Incucyte live-cell imaging system (Sartorius) normalized to vehicle treated organoids followed by normalization to *sgNTCi* after 10 days, cells were partially dissociated and related on day 7. Two-way ANOVA, multiple comparisons to *sgNTCi* within each dose. **b)** Immunoblot for indicated proteins in SNL tumour organoids after treatment with cisplatin from **Extended Data Fig. 8a**. **c)** Organoid growth after treatment with paclitaxel determined by GFP+ confluency using an Incucyte live-cell imaging system (Sartorius) normalized to vehicle treated organoids followed by normalization to *sgNTCi* after 5 days. Two-way ANOVA, multiple comparisons to *sgNTCi* within each dose. **d)** Human BCi-NS1.1 cell line growth after treatment with cisplatin for 48 hrs determined by confluency using an Incucyte live-cell imaging system (Sartorius) normalized to vehicle treated cells. **e)** Immunoblot for indicated proteins in human BCi-NS1.1 cell lines expressing non-targeting control (sgNTC) or sgKLF4 after 48 hours of treatment with the indicated doses of cisplatin. **f)** Human BCi-NS1.1 cell line growth after treatment with paclitaxel for 72 hrs determined by confluency using an Incucyte live-cell imaging system (Sartorius) normalized to vehicle treated cells. **For all panels,** error bars represent mean +/- SD; exact p-values are listed.

**Extended Data Fig. 9.**
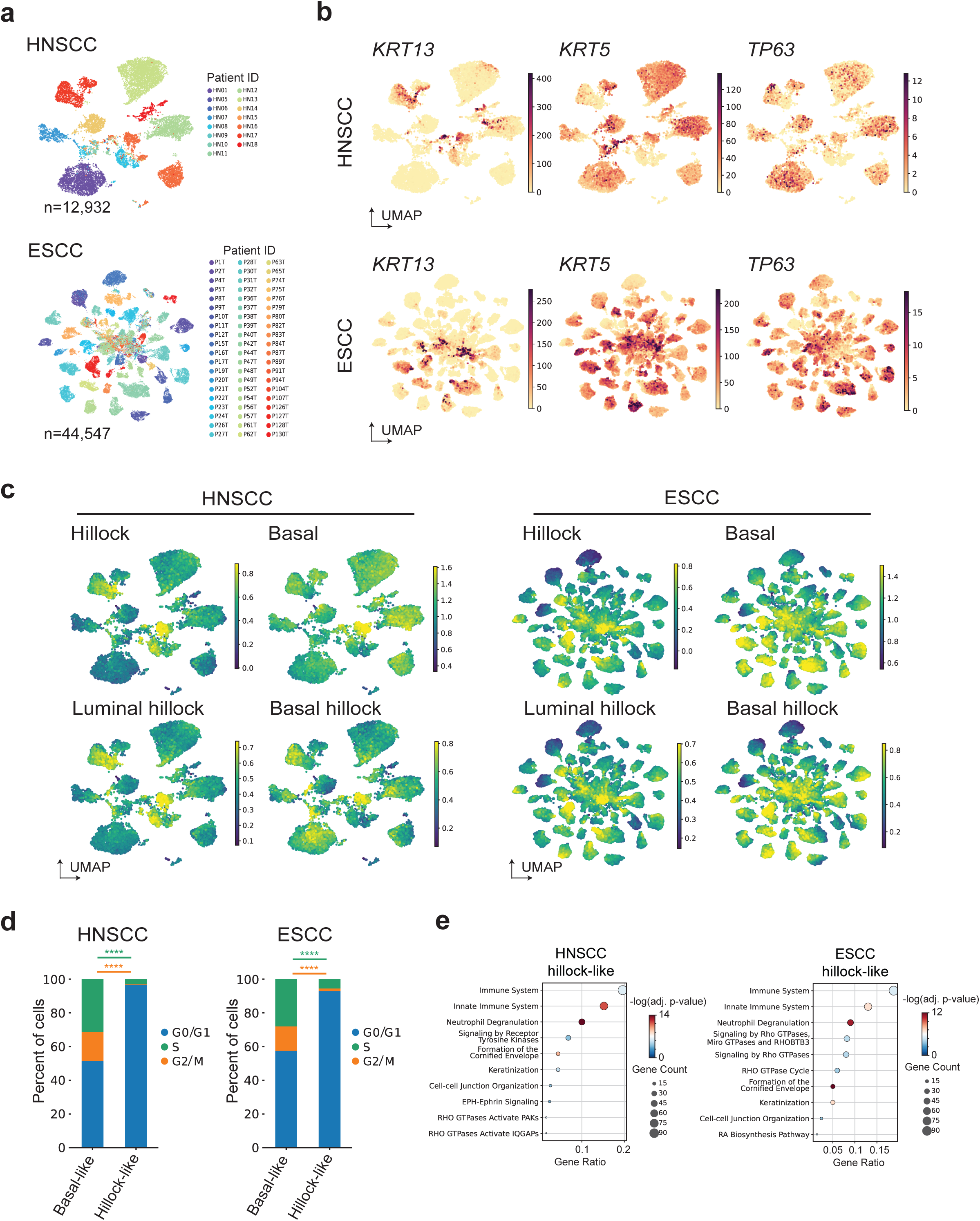
Hillock-like cells are present in squamous tumours of the esophagus and head & neck, related to Fig. 8. **a)** UMAP of scRNA-seq from human head & neck tumour cells from 15 patients^95^ or from esophageal tumours from 60 patients^96^ labeled by patient (HN, or P) sample ID. **b)** scRNA-seq expression of squamous (*TP63, KRT5)* or hillock markers (*KRT13*) in UMAP space from cells in **8a**. **c)** Human hillock score, basal score, luminal hillock score, and basal hillock score (**Supplementary Tables 1-5**) applied to tumour cells as in **8a** with score projected in UMAP. **d)** Cell cycle assignments applied to cells from **8a** (see **Methods**) shown as a stacked bar plot with percent of cells occupying each cell cycle phase by cell state assignment as in **8H**. Statistics shown compare G2/M (orange) and S-phase (green) gene scores between assigned cell states. Mann-Whiteney U-Test. ****p ≤ 0.0001. **e)** Gene set enrichment analysis comparing human tumour cells in **8b** by assigned cell type. The ENRICHR gene set used was ‘Reactome Pathways 2024’.

**Extended Data Fig. 10.**
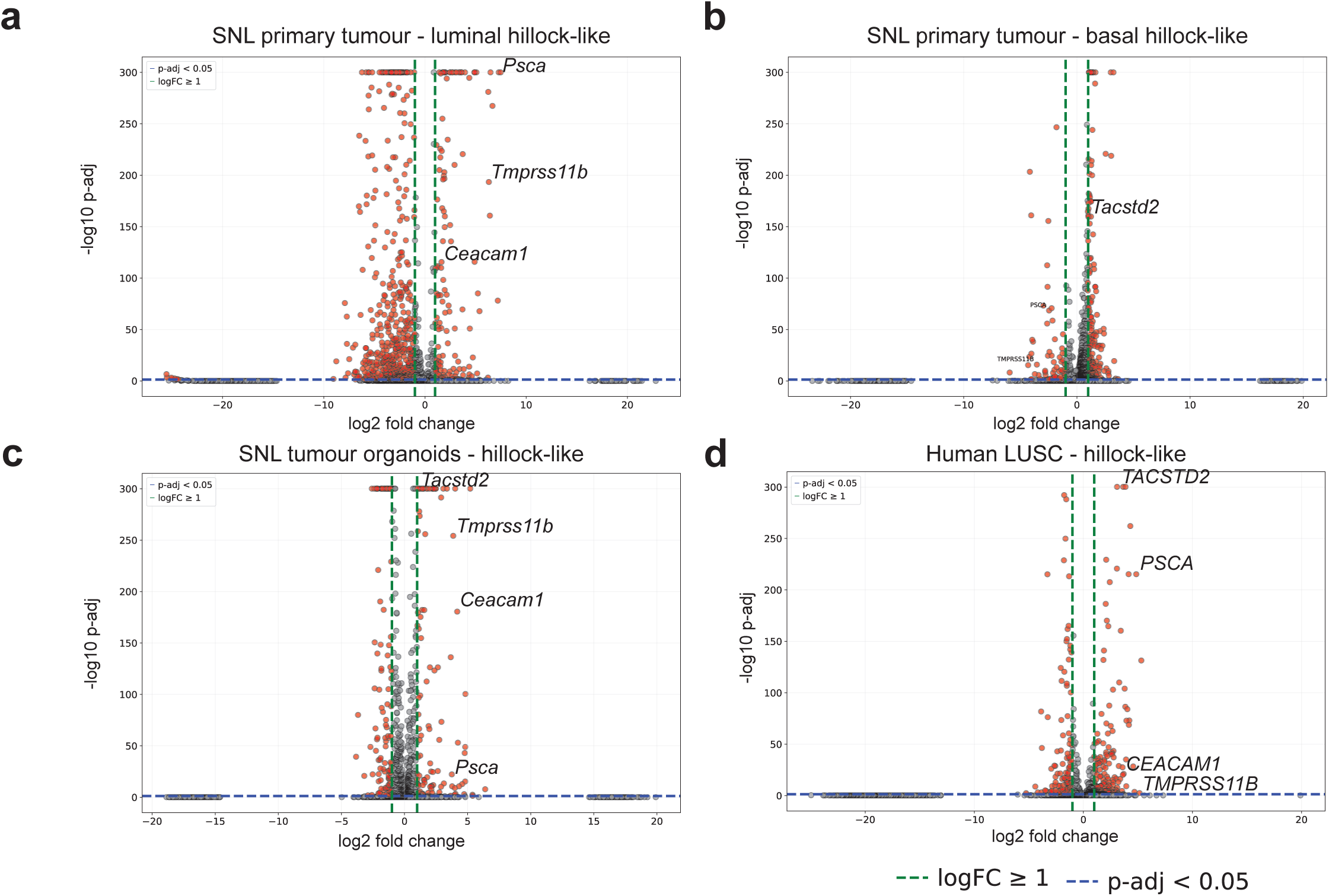
Differentially expressed cell surface protein-encoding genes by cell state using scRNA-seq data, related to Fig. 8. **a-d)** Volcano plots of DEGs by cell state filtered for predicted cell surface protein encoding genes. **See also Supplementary Table 15.**

